# The immunoglobulin A isotype of the Arabian camel (*Camelus dromedarius*) preserves the dualistic structure of unconventional single-domain and canonical heavy chains

**DOI:** 10.1101/2023.09.04.556198

**Authors:** Walter Conca, Soad Salih, Rana Al-Rabiah, Ranjit Parhar, Mahmoud Abd-Elnaeim, Hussein Al-Hindas, Alexander Tinson, Kate Collison, Uday Kishore, Futwan Al-Mohanna

## Abstract

Evolutionary processes of adaptive immunity in *Camelidae* and some cartilaginous fishes resulted in the concurrent expression of classic heavy-light chain heterotetrameric antibodies together with unconventional heavy chain-only homodimeric antibodies of the IgG class. Heavy chain-only IgG bears a single variable domain and lacks the constant heavy (C_H_) γ1 domain required for the covalent pairing with the light chain. It has so far not been reported whether this distinctive feature of IgG is also observed in the IgA isotype. Therefore, gene-specific primers were used to generate a library of IgA heavy chain cDNA derived from RNA extracted from the third eyelid of the dromedary where isolated lymphoid follicles and plasma cells abound at inductive and effector sites, respectively. Majority of the cDNA clones revealed the hallmarks of heavy chain-only antibodies, *i.e.* camelid-specific amino acid substitutions in framework region 1 and 2, broad length distribution of complementarity determining region 3, and the absence of the C_H_α1 domain. In a few clones, however, the cDNA of canonical IgA heavy chain was amplified which included the C_H_α1 domain, analogous to C_H_γ1 domain in dromedary IgG1 subclass. Moreover, we noticed a short, proline-rich hinge, similar to that in human IgA2, and, at the N-terminal end of the C_H_α3 domain, a unique, camelid-specific pentapeptide consisting of serine, glutamic acid, aspartic acid, alanine or threonine, and isoleucine, of undetermined function, designated as the inter-α region. Immunoblots using rabbit anti-camel IgA antibodies raised against C_H_α2 and C_H_α3 domains as well as the inter-α region revealed the expression of a ∼52 kDa and a ∼60 kDa IgA species, corresponding to unconventional and canonical IgA heavy chain, respectively, in tissue extracts of the third eyelid, *trachea*, small and large intestine. In contrast, the leporine anti-C_H_α1 antibody detected canonical, but not unconventional IgA heavy chain, in all the examined tissues, milk, and serum, in addition to another hitherto unexplored species of ∼45 kDa in milk and serum. Immunohistology using anti-C_H_α domain antibodies confirmed the expression of both variants of IgA heavy chains in plasma cells in the third eyelid’s lacrimal gland, and in the *lamina propria* of *conjunctiva*, bronchial and intestinal *mucosa*. We conclude that in the dromedary, the IgA isotype has expanded the immunoglobulin repertoire by co-expressing single-domain unconventional as well as canonical IgA heavy chains, comparable to the IgG class, thus underscoring the crucial role of heavy chain-only antibodies in immune defense not only in circulation but also at the mucosal frontiers.

## Introduction

Within the wider context of exploring mechanisms of adaptation of ocular immune defense to biohazards in the desert habitat, the present study was based on the fundamental knowledge that one-humped Arabian camels or dromedaries (*Camelus dromedarius*) share with other species of the *Camelidae* family, which includes *Camelus ferus* (two-humped wild or feral camel), *Camelus bactrianus* (two-humped Bactrian camel), *Lama glama* (llama), *Lama guanicoe* (guanaco), *Vicugna vicugna* (vicuña) and *Vicugna pacos* (alpaca), a distinguishing characteristic of circulating antibodies of the IgG isotype, namely the absence of the constant heavy (C_H_) γ1 domain that covalently pairs with the constant domain of the light chain (C_L_), resulting in the formation of homodimeric heavy chain-only (*alias* ‘only-heavy-chain’) antibodies where the paratope comprises the variable domain of the heavy chain variant, termed the variable heavy heavy domain (V_H_H/VHH/VH_H_ – all three acronyms are in use) (1–3). These unconventional antibodies were also discovered in the nurse shark (*Ginglymostoma cirratum*) and labeled as *neo* or nurse shark antigen receptor (4, 5). Over the years, these discoveries were widely exploited and resulted in several medical innovations (6–8). Although the etiology and pathways which drove B cell ontogeny in *Camelidae* to produce a functional IgG antibody alternative remain enigmatic, it is important to bear in mind that unconventional heavy chain-only antibody subclasses IgG2 (IgG2a and IgG2c) and IgG3 circulate beside canonical heterotetrameric subclass IgG1 (IgG1a and IgG1b), inferring that either antibody variant is a key effector in humoral immune defense (9–14). Whether in the Arabian camel other immunoglobulin classes share this exceptional feature of antibody structure has so far not been reported.

Therefore, we embarked on studies in the dromedary and focused on IgA as the established protagonist of adaptive immune defense at the ocular surface, which is protected by a tear fluid film produced by lacrimal and sebaceous glands as well as goblet cells in the *conjunctiva* (15–18). Primers deduced from amino acid sequences in constant domains of IgA heavy chain of alpaca and two-humped Bactrian camel served to create a library of IgA heavy chain cDNA amplified by PCR from mRNA isolated from the third eyelid of the dromedary, which harbors the *conjunctiva-*associated lymphoid tissue (CALT), a large sebaceous gland and the gland of Harder or Harderian gland (HG) (19–23). We show that most of the amplified IgA heavy chain transcripts were devoid of the C_H_α1 domain, the fingerprint of heavy chain-only antibodies, while few corresponded to conventional IgA heavy chain transcripts. We also provide protein data confirming the translation of both IgA subclasses not only in CALT, but also in the upper respiratory and the intestinal tract. These observations fill a gap in our knowledge about the composition of the dromedary’s mucosal IgA, and favor an evolutionary scenario in which exposure to microbial pathogens and commensal microflora (24), probably forced by environmental hazards in the desert habitat, such as unabated exposure to UV light, temperature and humidity extremes (25), as well as metabolic and osmotic derangements during starvation and dehydration (26), accounted for a distinct and likely “rescue” or “salvage” pathway of antibody synthesis capable of complementing or alternating with, yet not replacing, the highly complex, fine-tuned endoplasmic assembly of heavy and light chains (27–29).

## Materials and methods

### Animal studies

Third eyelids *in toto,* tissue samples of the *trachea*, as well as the small and large intestine were harvested at necropsy from three apparently healthy, adult male one-humped camels (C1, C2 and C3) at the municipal abattoir in Riyadh, immediately fixed in 10% (v/v) neutral buffered formalin for >24 hours for histomorphology and immunohistochemistry, or snap frozen in liquid nitrogen for RNA and protein extraction. After fixation, tissues were paraffin-embedded and cut into 4-μm sections. Sequential sections were either stained with hematoxylin and eosin (H&E), some with Masson’s trichrome stain, or analyzed by immunohistochemistry. Tissues of camel C3 were used for further analyses. Blood was drawn by venipuncture from the external jugular vein from healthy female and male dromedaries at a camel farm in Buraidah, Central Region of Saudia Arabia, allowed to clot and *sera* were stored at −20°C. Camel milk was collected at a local farm in the vicinity of Riyadh, Central Region of Saudi Arabia. Approval to carry out the sampling procedures was obtained from “The Animal Care and Use Committee” of the King Faisal Specialist Hospital & Research Center (RAC#2180-023).

### RNA extraction, rapid amplification of cDNA ends (RACE) and cDNA library construction

Total RNA was isolated from the third eyelid of dromedary C3 with Zymo® according to the manufacturer’s instructions (Zymo Research, Irvine, CA, USA). RNA was quantified by NanoDrop™ spectrophotometer (Thermo Fisher Scientific, Waltham, MA, USA) and its integrity assessed by 2100 Bioanalyzer instrument (Agilent Technologies, Santa Clara, CA, USA). We first generated a ∼500 base pair (bp) product using primers derived from the constant domains of IgA heavy chain from alpaca and Bactrian camel. The cDNA sequences obtained showed ∼97% identity with nucleotide sequences of constant domains of IgA heavy chains of these two camelids. Thereafter, we amplified further towards the N-terminal and C-terminal end of the cDNA using the RACE technique. RACE is a PCR-based technique which facilitates the cloning of full-length cDNA sequences. RACE is widely used to amplify unknown cDNA sequences towards the 5’ and 3’ ends of mRNA using gene-specific primers (GSP) derived from a known short internal stretch of the mRNA of interest (30). RACE was performed with the Invitrogen™ GeneRacer™ Kit (Thermo Fisher Scientific, Waltham, MA, USA) to generate full-length cDNA as follows: 4μg of total RNA was dephosphorylated with calf intestine alkaline phosphatase to remove free 5’ phosphates from rRNA, fragmented RNA, tRNA and contaminating DNA. Full-length capped mRNA was treated with tobacco acid pyrophosphatase to remove the 5’ cap structure. This treatment left a phosphate at the 5’ end required for ligation to 44-base-long GeneRacer™ RNA Oligo (5’- CGACTGGAGCACGAGGACACTGACATGGACTGAAGAGTAGAAA-3’) using T4 RNA ligase, which provides a priming site for GeneRacer™ 5’ primer and 5’ nested primer. Ligated mRNA was reverse transcribed using Superscript™ III Reverse Transcription (RT) and the 60-base GeneRacer™ oligo (dT) primer (5’- GCTGTCAACGATACGCTACGTAACGGCATGACAGTG(T)_24_-3’) containing a dT tail of 24 nucleotides to create RACE-ready first-strand cDNA with known priming sites at the 5’ and 3’ ends, and restriction sites for efficient cDNA synthesis for cloning and sequencing purposes after PCR amplification. The GeneRacer™ oligo (dT) primer sequence contains the priming sites for the GeneRacer™ 3’ and the GeneRacer™ 3’ nested primers. After comparing nucleotide sequences of constant domains of IgA heavy chains of other camelids, reverse (5’- AGCAGGTGGACCTGGGGCCGGAT-3’) and forward (5’- ATCCGGCCCCAGGTCCACCTGCT-3’) GSP were chosen from the C_H_α3 domain of IgA heavy chain of the Bactrian camel (GenBank #KP999944.1; nucleotides 643-665 of a partial coding sequence; N-*terminus* of C_H_α3 domain) using the online primer designing tool in NCBI (NCBI Primer Blast) to amplify the 5’ and 3’ ends of dromedary IgA heavy chain cDNA, respectively. As an internal control, *β-actin* mRNA was amplified. PCR amplifications of 1 μl RT template were performed in 50 μl aqueous reaction mixture consisting of High-Fidelity PCR buffer, 2 mM MgSO_4_, 200 nM of each dNTP, 200 nM forward or reverse GSP, 600 nM GeneRacer™ 5’ primer or 3’ primer, and 2.5 U Platinum® *Taq* DNA polymerase High Fidelity that generated blunt end products (Thermo Fisher Scientific, USA), using hot start and touchdown PCR, following the manufacturer’s recommendations. To obtain 5’ ends, first-strand cDNA was amplified using the reverse GSP and the GeneRacer™ 5’ primer and 5’ nested primer. Only mRNA that had the GeneRacer™ RNA Oligo ligated to the 5’ end and was completely reverse transcribed was amplified by PCR. To obtain the 3’ ends, first-strand cDNA was amplified using the forward GSP and GeneRacer™ 3’ primer and GeneRacer™ 3’ nested primer. Only mRNA that had a poly(A) tail and was reverse transcribed was amplified by PCR. Three negative PCR controls (no template; no GSP; no GeneRacer™ 5’ or 3’ primers) were included to eliminate non-specific amplification. Cycling parameters were as follows: 1 cycle: 2 min at 94°C; 5 cycles: 30 sec at 94°C and 1min/1kb DNA at 72°C; 25 cycles: 30 sec at 94°C; 45 sec at 65°C and 2 min at 68°C; 1 cycle for 10 min at 70°C. After PCR, 10 μl of the amplified product was analyzed on a 1.2% (w/v) agarose gel.

### Cloning and sequencing

PCR-amplified 5’ and 3’ cDNA fragments were separated on a 1.2% (w/v) agarose gel. Bands were eluted using QIAquick Gel Extraction Kit (Qiagen, Inc., Chatsworth, CA, USA), then ligated into Zero Blunt™ TOPO™ vector (Invitrogen, Carlsbad, CA, USA) using DNA Ligation Kit, Mighty Mix (Takara Bio USA, San Jose, CA, USA), and transformed into chemically competent TOP10 *E.coli* cells (Invitrogen, Carlsbad, CA, USA). Individual colonies were grown, and plasmids purified using a QIAprep Spin Miniprep Kit (Qiagen, Inc., Chatsworth, CA, USA). For analysis of the 5’ RACE products, those colonies with complete sequence of the GeneRacer™ RNA Oligo and reverse GSP were selected for further analysis, indicating that the intact 5’ RACE product was ligated to GeneRacer™ RNA Oligo. DNA was isolated from selected transformants and sequenced from both directions using M13 forward and M13 reverse sequencing primers with BigDye™ Terminator v3.1 Cycle Sequencing Kit (Applied Biosystems™, Inc. Foster City, CA, USA) at the King Faisal Specialist Hospital & Research Center sequencing facility. Forward and reverse sequences were aligned using EMBOSS needle pairwise sequence alignment algorithms and DNASTAR™ program and merged using EMBOSS. The same strategy was used for 3’ RACE colonies obtained after amplification in the presence of GeneRacer™ oligo (dT) primer sequence and forward GSP. Overlapping 5’ and 3’ RACE sequences were then combined, and the complete cDNA sequence of IgA heavy chain obtained.

### Generation of rabbit polyclonal antibodies against constant domains of dromedary IgA heavy chains and the inter-α region

From the following three conceptually translated amino acid sequences of the C_H_α1, C_H_α2 and C_H_α3 domains the cDNA sequence was derived, His tagged and expressed in *E. coli*:

1. C_H_α1 SPSVFPLGPSYDKASRQVALACLVHGFFPPAPLKVTWGLSGQNVSVMDFPAVQPASGVLYTMSSQLTTPVEQCPDSEIVTCQVQHLSSSSKTVNVPCK
2. C_H_α2 CCNPSLALHPPALEDLLLGSNASLTCTLSGLRNPEGAQFTWTPSGGKVAVQQSPKSDPCGCFSVSSVLPGCAEQWNSKTNFSCSATHPESKNTLYATITK
3. C_H_α3 IRPQVHLLPPPSEELALNEMVTLTCVVRGFSPKDVLVRWLHGNQELPREKYLTWRPLPEPEQSITTYAVTSLLRVEAEAWKQGDNYSCMVGHEALPLAFTQKTIDRLSGKPTHVNVSVVMAEAEGVCY

All proteins were custom-made by GenScript HK Limited, Hong Kong SAR (PolyExpress Premium antigen specific affinity purified pAb package). Freund’s adjuvant was added to the immunogen. Six New Zealand rabbits (two rabbits/recombinant protein) were immunized with three injections by conventional protocol. Seven days after the third immunization, the titer of antiserum was tested by ELISA to detect antibody response. Total IgG from pre-immune serum was used as a negative control. Rabbits were sacrificed according to animal welfare principles. The final antiserum was purified by antigen affinity column. 0.02% (w/v) sodium azide was added as preservative, and antibody preparations were stored at −20°C.

The oligopeptide, ATITKSSEDAIRPQVC, spanning the inter-α region (IAR), was generated with a cysteine added at the C-*terminus* by 9-fluorenylmethyloxycarbonyl (Fmoc) solid phase peptide synthesis on resin and then removed from the resin by treatment with trifluoroacetic acid (31). Oligopeptides were conjugated with keyhole limpet hemocyanin, then purified by reverse-phase HPLC with acetonitrile and water and verified by mass spectrometry. The purified product was used for immunization of two rabbits following the same protocol adopted for the immunization with C_H_α domains and the antibody was purified by antigen affinity column.

### Protein preparation, gel separation and immunoblotting

Serum lysis was done in ice-cold RIPA buffer (Sigma-Aldrich, St. Louis, MO, USA) containing cOmplete™ Mini protease inhibitors cocktail (Roche Diagnostics, Mannheim, Germany). Similarly, RIPA lysates were made from snap-frozen samples of various camel tissues. 10-20 µg of total protein were mixed with SDS-PAGE loading buffer, pH 7.0, containing 5% (v/v) β-mercaptoethanol, heated to 95°C for 5 min, and separated by SDS-PAGE at 20 mA using NuPAGE™ 10% (w/v), Bis-Tris, 1.5 mm, Mini Protein gels (Invitrogen, Carlsbad, CA, USA) in a Mini-PROTEAN Tetra vertical electrophoresis cell (Bio-Rad, Hercules, CA, USA). The gels were fixed and stained with 50% (v/v) methanol, 10% (v/v) acetic acid, 0.1% (w/v) Coomassie Brilliant Blue R-250, and de-stained with 40% (v/v) methanol and 10% (v/v) acetic acid. Proteins were then transferred to 0.2 µm nitrocellulose membranes (Bio-Rad, Hercules, CA, USA) using the Trans-Blot® Turbo™ semi-dry transfer system (BioRad, Hercules, CA, USA) Anti-C_H_α1 domain (1/2500), anti-C_H_α2 domain (1/2500), anti-C_H_α3 domain (1/2500) and anti-IAR (1/2500) primary antibodies, diluted in 3% (w/v) non-fat dry milk, were added in TBS-Tween buffer overnight at 4°C, followed by washing in TBS-Tween buffer. Thereafter, horseradish peroxidase (HRP)-conjugated goat anti-rabbit IgG antibody (1:10000) (Promega, Madison, WI, USA), diluted in 3% (w/v) non-fat dry milk, was added and the blot incubated in TBS-Tween buffer for 2 hours at room temperature, followed by three washes with PBS-Tween buffer. Proteins were detected using Pierce™ ECL Plus Western Blotting Substrate (Thermo Fisher Scientific, Waltham, MA, USA) and images were acquired via ChemiDoc Imaging System (Bio-Rad, Hercules, CA, USA). Densitometry analysis was performed using Image Lab software (Bio-Rad, Hercules, CA, USA).

### Immunohistochemistry

*ultra*View® Universal DAB Detection Kit (Ventana Medical Systems, Oro Valley, AZ, USA) in combination with BenchMark ULTRA IHC/ISH System (Ventana Medical Systems, Oro Valley, AZ, USA) fully automated instrument, was used for the detection of IgA heavy chains in formalin-fixed paraffinized tissue sections of various organs using rabbit anti-C_H_α1, C_H_α2, and anti-IAR primary antibodies. The reagents provided with the kit do not contain biotin, and therefore non-specific staining from endogenous biotin is eliminated. Deparaffinization was done at 72°C, followed by cell conditioning (heat-induced epitope retrieval) with ULTRA CC1 solution for 64 min at 95°C. Primary antibodies were applied manually, all other reagents automatically in a pre-diluted dispenser to tissue sections placed on glass microscope slides and were then exposed to a “cocktail” of secondary antibodies, consisting of goat anti-rabbit IgG, goat anti-mouse IgG and IgM, conjugated to HRP by long-arm linkers (HRP multimer). Primary antibody incubation time was 20 min, followed by secondary antibody incubation time for 8 min at room temperature. The immune complex was then visualized with the chromogen 3,3’-diaminobenzidine tetrahydrochloride (DAB) and H_2_O_2_, which upon oxidation formed a brown precipitate. Secondary antibody alone was used as a negative control. Counterstaining was performed with hematoxylin first, followed by post-counterstaining with bluing reagent for 8 min, respectively. Microphotographs were taken with an Olympus BX53 digital camera (Olympus Corporation, Tokyo, Japan).

## Results

### Anatomy and histomorphology of the dromedary’s third eyelid

Gross anatomical details of the eye, skull and third eyelid are shown in Fig 1. An image of the left eye shows details of the eyebrow, the upper and lower eyelids with long eyelashes, and the eyeball (*bulbus oculi* or globe) with hyperpigmentation of the sclera in the sun-exposed segment. The semilunar fold with a dark-brown rim at the rostro-medial angle (white arrow) belongs to the retracted third eyelid (*alias* nictitating membrane or *palpebra tertia*), which on blinking unfolds across the melanin-rich globe in a posterior direction, asynchronously to the orthogonal movement of the upper and lower eyelids (Fig 1A). An anterior-posterior view of the skull demonstrates the orbital bones sticking out in a lateral and inferior orientation (Fig 1B). Notably, massive orbital roofs with a deep fissure-like supraorbital notch provide shade from the incessant sun exposure and protection from sand grains during windy weather (black block arrows). A pair of third eyelids excised *in toto* is shown (Fig 1C, D). They have the shape of a trapezoid, with a slightly convex, smooth, and hence shiny, melanin-rich palpebral (outer) (Fig 1C) and a concave, rough and less pigmented bulbar (inner) surface (Fig 1D). The base of the third eyelid is wider (∼3 cm) and thicker than the shorter (∼1.5 cm) and thinner, hyperpigmented free edge (red arrows). In the center, a long cartilage shaft traverses the body of the nictitating membrane from its root at the base and ramifies, Y-shaped, into two terminal branches (blue arrows). At the base of the third eyelid, we observed two large, structurally and functionally different glands next to each other. As shown in Fig 1C, the first and more superficial sebaceous gland (SG), topographically and functionally corresponding to the caruncle in humans, consists of numerous yellowish-beige lobules. The second and deeply located gland is the compact, lobulated, oval-shaped, pink-brownish mucous lacrimal gland, best known as the HG (black arrows). Here, most of the proteins found in the middle aqueous layer of tear film are synthetized, including secretory IgA (sIgA), as in other species (22, 32), whereas the SG is the principal factory of lipids that constitute the insulating outermost lipid layer of tear film (33). The innermost layer of tear film consists mainly of mucins secreted by goblet cells located in the *epithelia* of the glandular excretory ducts and the *conjunctiva* (34).

**Figure 1.**
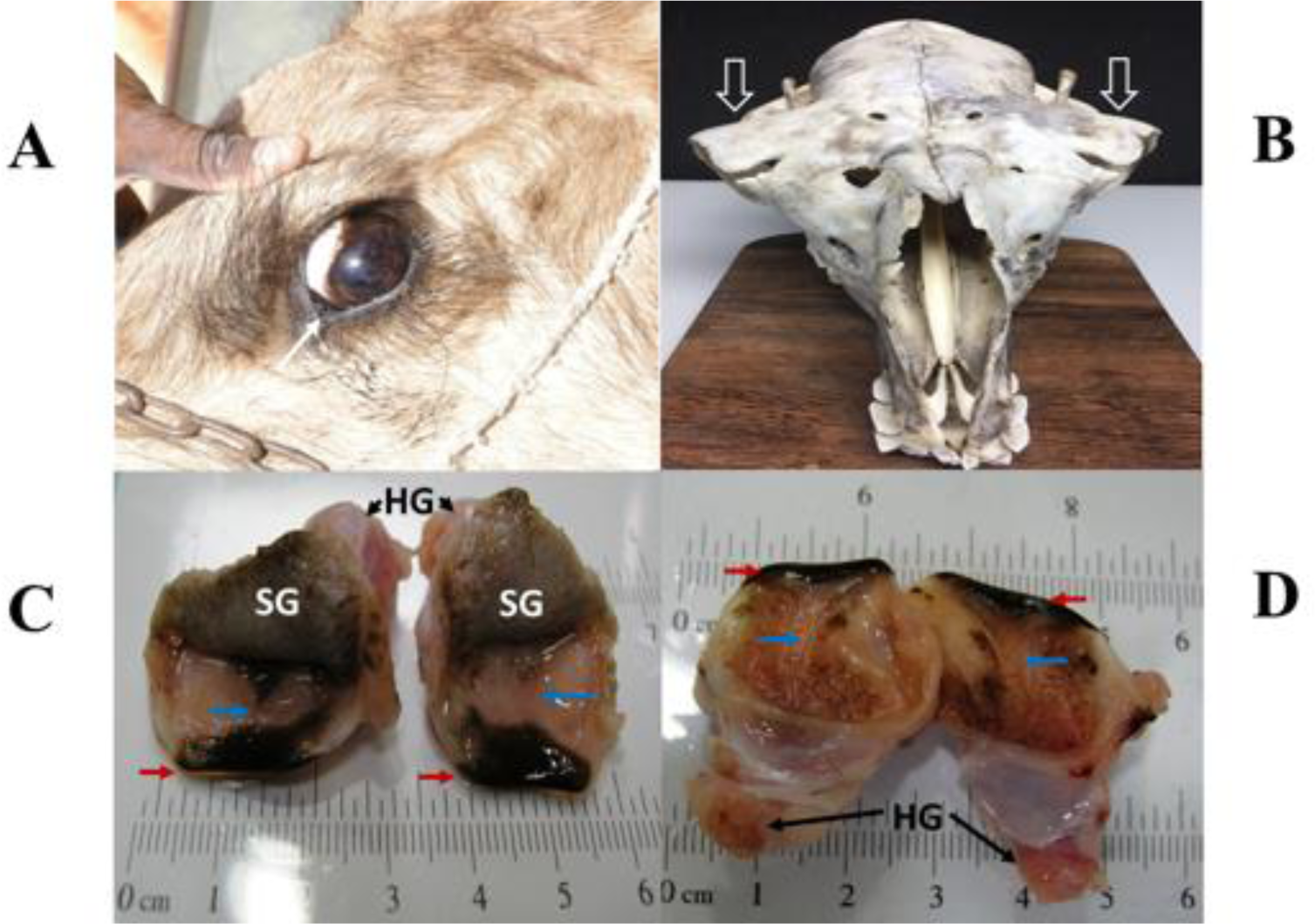
Eye, skull and third eyelid of the dromedary. In (A), a close-up image of the left eye of a dromedary, with eyebrow and upper eyelid raised, reveals long eyelashes, the melanin-enriched *sclera* in its sun-exposed portion, and the hyperpigmented free margin (white arrow) of the retracted third eyelid. In (B), an anterior-posterior view of the flat skull demonstrates remarkably protruding *orbitae* (black block arrows), and behind these, the condylar processes of the mandible sticking out from the temporal *fossae*. In (C), the palpebral (outer) surfaces, and in (D) the bulbar (inner) surfaces of an excised pair of third eyelids are shown. Third eyelids are trapezoidal in shape, have melanin-enriched free margins (red arrows) and are supported by Y-shaped cartilage shafts (blue arrows). Two large glands are incorporated in the third eyelid: the superficial sebaceous gland (SG), and the deep Harderian (lacrimal) gland (HG) (black arrows).

The salient histomorphologic features of the third eyelid are shown in Fig 2. In Fig 2A, the sequential arrangement of the principal glands of the third eyelid is shown in a single cut. The superficial lobulated SG is located anterior (left) to the conjunctival *fornix* (*), a deep reflection of the *conjunctiva*, where the main excretory ducts of the HG are lined with many large goblet cells (arrow). Lining the *fornix*, the *conjunctiva* consists of simple columnar *epithelium*. Posterior (right) to the *fornix*, the palpebral part of the lobulated HG reveals the compact structure of a tubulo-alveolar mucous gland. As shown in Fig 2B, the bulbar part of the HG is adjacent to an agglomerate of isolated lymphoid follicles (ILF) lacking germinal centers, each one situated beneath the specialized follicle-associated *epithelium* (arrow) and the subepithelial dome (*), a complex functional unit that consists of a cluster of B cells built on stromal cells, surrounded by many dendritic cells and few T cells. Here, at the conjunctival cul-de-sac, antigen uptake *via* M cells and processing by antigen-presenting cells is known to take place, followed by IgA class switching in B cells, mostly independent of T cells. Therefore, ILFs represent an inductive site for conjunctival IgA antibody responses to airborne antigens at the ocular surface and resemble the architecture of lymphoid tissue in *mucosa*-associated lymphoid tissues (MALT) of other species (35–43). In Fig 2C, the central part of the HG is shown to enclose the root of the hyaline cartilage (C) covered by *perichondrium* (PC). In Fig 2D, *acini* in the secretory endpieces of the HG are lined with columnar cells with basally situated nuclei and a narrow *lumen* (*). They are surrounded by *stroma* which harbors the effector site of the humoral immune response as it is enriched with plasma cells (arrows). The lobules of the branched tubulo-alveolar holocrine SG consist of *acini* without *lumen* (Fig 2E). The terminal differentiation of sebocytes (S) is accompanied by the progressive accumulation of lipid droplets, morphologically resembling those in multilocular brown adipocytes, and ultimately results in *karyopyknosis* (arrow) with the release of the lipid contents and nuclear material (purple by Masson’s trichrome staining) into duct-like conjunctival invaginations (*). In the center of this photomicrograph, a pilosebaceous unit was observed wherein a small sebaceous gland is found adjacent to a hair shaft (H; red). The molecular processes underlying this unique type of programmed cell death of sebocytes, which we propose to name “seboptosis”, were studied in detail in the SG of the murine *praeputium* and found to be associated with the breakdown of lysosomes, release of lysosomal DNase2 into the cytoplasm, nuclear degradation, and subsequent complete disintegration of sebocytes (44). In Fig 2F, a section through the body of the third eyelid, parallel to the long axis of the tapering cartilage shaft (C) covered by *perichondrium* and surrounded by a thick layer of dense connective tissue (CT), shows the stratified squamous *epithelium* of the *conjunctiva* (Conj) on both sides and higher vascularity (V) on the palpebral side (pConj) with abundant intraepithelial pigmented granules (oblique arrows). By contrast, on the bulbar side, no granules were observed (bConj). Lymphoid cells are found sparse throughout the subepithelial layers.

**Figure 2.**
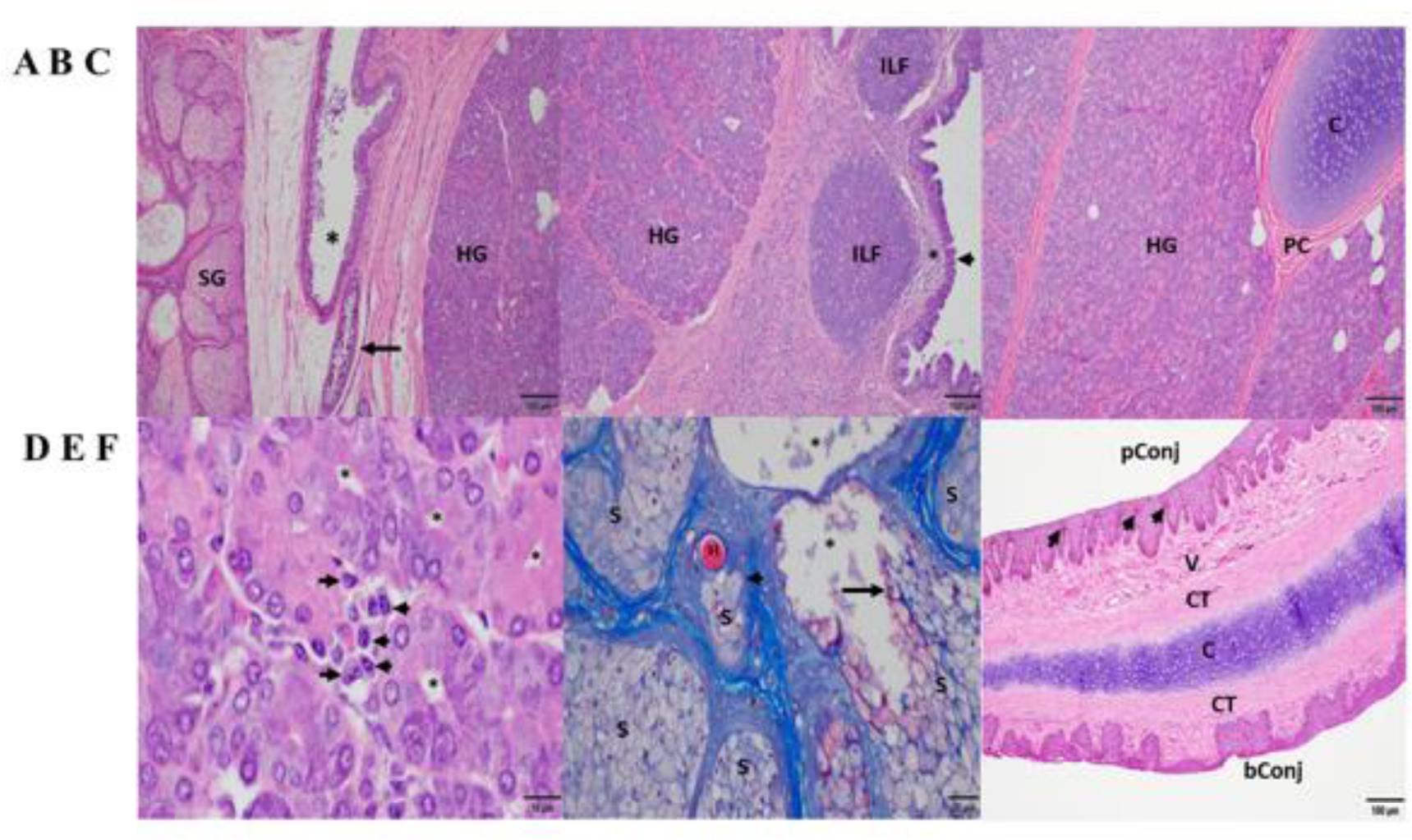
Histomorphology of the third eyelid of the dromedary. In (A), the superficial, holocrine sebaceous gland (SG) and the deep, merocrine Harderian (lacrimal) gland (HG) are separated by the deep *fornix* (*) of the *conjunctiva* in the center, where a goblet cell-enriched main duct (arrow) collects the lacrimal fluid. In (B), posterior (bulbar) to the HG, numerous isolated lymphoid follicles (ILF) are found beneath subepithelial domes (*) and the specialized follicle-associated *epithelium* (arrow) of the *conjunctiva*. In (C), the root of the hyaline cartilage (C) covered by *perichondrium* (PC) is embedded in the lobulated and septated HG, with a few interspersed white adipocytes. In (D), numerous plasma cells (arrows) cluster near *acini* with a central *lumen* (*). In (E), *acini* of the SG in contrast have no *lumen* and are packed with sebocytes (S), which upon terminal differentiation undergo *karyopyknosis* (long arrow; purple discoloration), a form of programmed cell death (“seboptosis”), followed by the release of lipid droplets and cellular debris (including nuclear material) into duct-like conjunctival invaginations (*). The red-stained hair shaft (H) is part of a pilosebaceous unit (short arrow). In (F), a section along the long axis of the body of the third eyelid shows the tapering hyaline cartilage shaft (C), encased by *perichondrium* and a dense layer of connective tissue (CT). Keratinocytes in the palpebral (outer) *conjunctiva* (pConj) contain pigmented granules (arrows). Many small vessels (V) are found in the subepithelial layer on the palpebral side, contrasting the scarcity of vasculature underneath the bulbar (inner) *conjunctiva* (bConj). Magnifications and scale bar lengths are as follows: in (A), (B), (C), and (F), 10x and 100 μm, respectively; in (D), 100x and 10 μm, respectively; in (E), 40x and 20 μm, respectively. Sections shown in (A), (B), (C), (D) and (F) were stained with H&E, section (E) was stained with Masson’s trichrome.

### Amplification products obtained via 5’ and 3’ rapid amplification of cDNA ends (RACE) and sequence analysis

5’ RACE PCR products separated on 1.2% (w/v) agarose gels revealed two species: most were ∼800 bp, a few ∼1100 bp long (Fig 3A, left panel). In contrast, a single amplification product of ∼450 bp length was obtained from 3’ RACE (Fig 3A, right panel). 5’ and 3’ PCR species were ligated into Zero Blunt™TOPO™ vector for sequencing. Out of 124 clones analyzed, two corresponded to the longer 5’ PCR products and included the C_H_α1 domain, whereas all other clones of the shorter 5’ PCR product lacked the C_H_α1 domain, which defines an unconventional IgA heavy chain. 5’ and 3’ RACE PCR products were joined together through their overlapping ends to generate two species of full-length cDNA of ∼1.3 kbp and ∼1.6 kbp, corresponding to unconventional and canonical IgA heavy chain, respectively. For comparative purposes, representative, conceptually translated complete protein sequences of classic IgA heavy chain (IgA classic; 485 amino acids; top) and unconventional IgA heavy chain (IgA HC; 388 amino acids; bottom) are shown in Fig 3B. In the center of C_H_α1 domain, starting from an internal methionine, we aligned another amino acid sequence (IgA*classic; 292 amino acids; middle), the partial sequence of “IgA alpha-1 constant region” (UniProt #A0A5N4E0V1. A0A5N4E0V1_CAMDR) translated from the whole genome shotgun sequence “Camelus dromedarius bred African isolate Drom800 chromosome 6/IgA alpha-1 constant region” (GenBank®# JWIN03000006.1; protein_id #KAB1277053.1) (45). Individual components of the variable domain of classic IgA heavy chain (V_H_) and unconventional IgA heavy chain (V_H_H), *i.e.* framework regions (FR) 1, 2, 3 and 4, and complementarity determining regions (CDR) 1, 2 and 3, as well as constant domains C_H_α1, C_H_α2 and C_H_α3, are indicated above the amino acid sequence, with normal and bold font letters alternating for a better illustration. Variable domains were defined according to IMGT^®^ delimitations (http://www.imgt.org, IMGT Scientific chart>Numbering>IMGT unique numbering for V-DOMAIN and V-LIKE DOMAIN) (46). A 19-mer leader peptide precedes the constituents of the variable domain FR1, CDR1, FR2, CDR2, FR3, CDR3 and FR4. The multiple constant domains of classic IgA heavy chain are numbered from the N-terminal to the C-terminal end as follows: C_H_α1, C_H_α2 and C_H_α3. The proline-rich (VPPPPPP) hinge region is positioned between the C_H_α1 and the C_H_α2 domain in canonical IgA heavy chain, which like its unconventional counterpart ends in the tailpiece (Tp), an 18-mer oligopeptide extension critical for the formation of IgA polymers. These constant domains were specified according to the scheme of human myeloma IgA1 Bur and IgA2 But (47, 48). In contrast, in unconventional IgA heavy chain, the C_H_α1 domain spanning 102 amino acids is absent, as indicated by dashes, and therefore, FR4 of the V_H_H domain, with the canonical J motif W-G-X-G and the variable/constant switch region consisting of the tripeptide VSS, directly precedes the heptameric hinge, which was identical in all clones. Downstream of the hinge region, we observed 100% identity among the variants of dromedary IgA heavy chains, except for T in place of A (underlined) at the N-*terminus* of C_H_α3 in IgA*classic heavy chain. 119 amino acid sequences of heavy chain-only IgA were aligned in the Supplementary file #1. In Fig 3B, key substitutions with hydrophilic amino acids in FR1 (S for L) and FR2 (F for V, E for G, R for L, G for W) of heavy chain-only IgA are underlined as they are considered crucial for the conformation and solubility of the V_H_H domain (2). As expected, the highest variability in length and amino acid composition was observed in the V-D-J recombined CDR3, ranging from 14 to 26 amino acids in our library. Of note, we observed a distinct pentapeptide, consisting of S, E, D, A (or T) and I at the N-terminal end of the C_H_α3 domain, intriguingly different from other mammals at this site (Fig 4). We labeled the pentapeptide as the IAR and used it to immunize rabbits.

**Figure 3.**
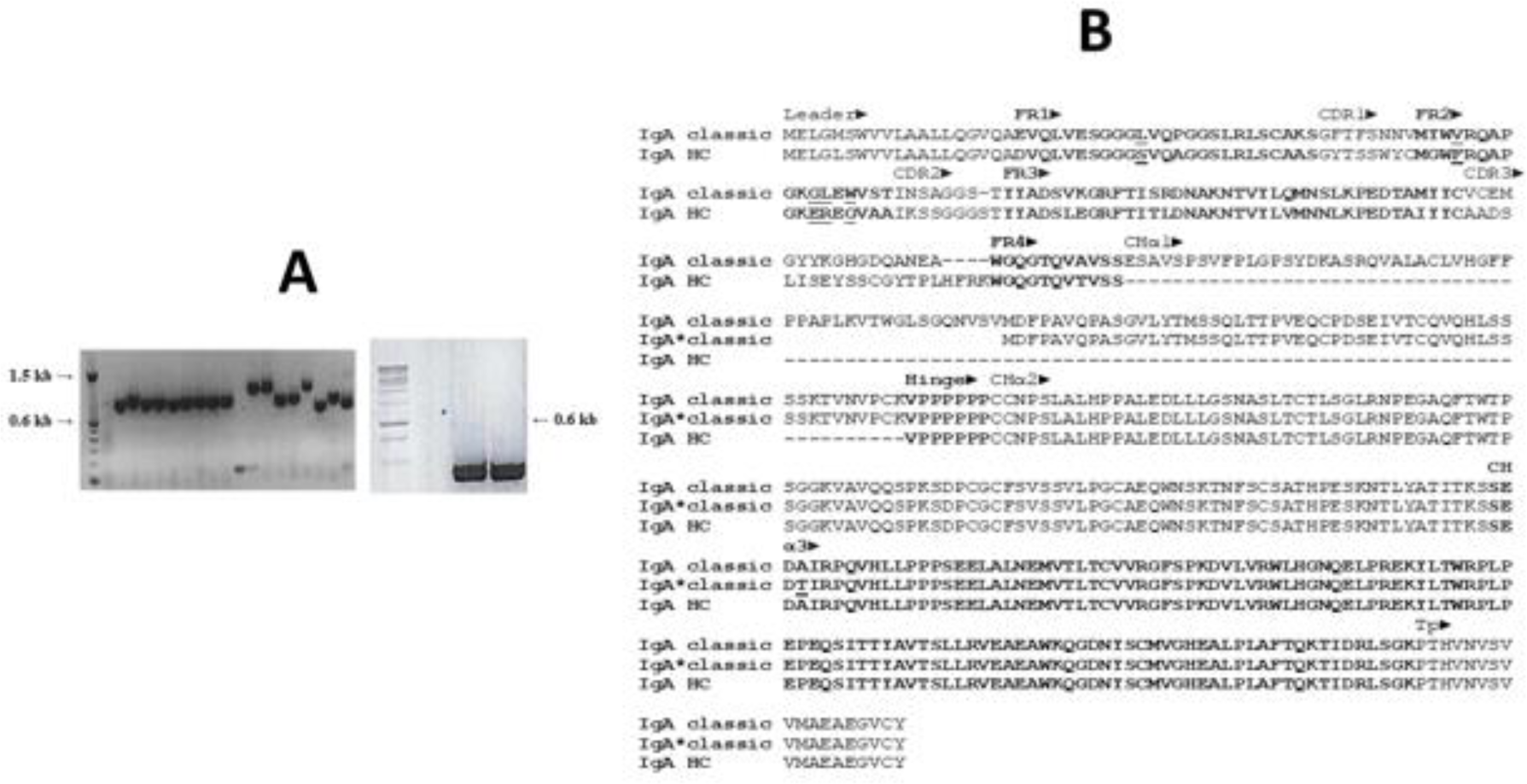
Rapid amplification of 5’ and 3’ cDNA ends (RACE) of dromedary IgA heavy chain and primary structure of classic and unconventional IgA heavy chain. In (A), PCR products of 5’ RACE (left panel) and 3’ RACE (right panel) are shown after agarose gel electrophoresis, with a DNA ladder loaded in the first well. The majority of 5’ PCR species were ∼800 bp, a few were ∼1.1 kbp. In contrast, 3’ RACE PCR species consisted uniformly of ∼450 bp. There were some minor variations in the length of the ∼800 kbp 5’ PCR products, reflecting the heterogeneity of CDR in V_H_H domains. The difference of ∼300 bp among the two 5’ PCR species, however, was explained by identification of the C_H_α1 domain in classic IgA heavy chain. In (B), two representative amino acid sequences of dromedary IgA heavy chains chosen from our library were aligned, with conventional IgA heavy chain (IgA classic) at the top and unconventional IgA heavy chain (IgA HC) at the bottom. We added a middle row (IgA*classic), a partial amino acid sequence with 292 residues of “IgA alpha-1 chain C region” translated from genomic DNA of the dromedary (45). This conceptually translated genomic sequence (IgA* classic) was incomplete as it did not extend beyond an internal methionine in the center of the C_H_α1 domain, therefore excluding 50 amino acid residues of the C_H_α1 domain upstream and the entire V_H_ domain. Variable domains of both heavy chains (V_H_ and V_H_H), preceded by a 19-mer leader peptide, comprised four framework regions (FR1, FR2, FR3 and FR4) and three complementarity determining regions (CDR1, CDR2 and CDR3), whereas constant domains of classic IgA heavy chain encompassed three (C_H_α1, C_H_α2 and C_H_α3), in contrast to only two domains (C_H_α2 and C_H_α3) in the unconventional format. A short hinge region, consisting of valine and six consecutive prolines, was located between C_H_α1 and C_H_α2 domains in conventional IgA heavy chain, in contrast to the hinge region in heavy chain-only IgA which separated FR4 of the V_H_H domain from the C_H_α2 domain. The tailpiece (Tp) represented the C-terminal end of all IgA heavy chains variants. Dashes indicate deficient single amino acid residues as well as lack of the entire C_H_α1 domain in heavy chain-only IgA. Definition of the constituents of the variable and constant domains (marked by alternating normal font and bold letters) was accomplished according to IMGT^®^ delimitations (46) and the numbering scheme of human IgA1 and IgA2 chains, respectively (47, 48). Underlined residues in FR1 and FR2 indicate amino acid changes considered critical for conformation and solubility of the V_H_H domain of heavy chain-only IgA. Further downstream, a unique pentapeptide (SEDAI) marked the N-*terminus* of the C_H_α3 domain, with threonine substituted for alanine in the whole genome shotgun IgA* classic sequence.

**Figure 4.**
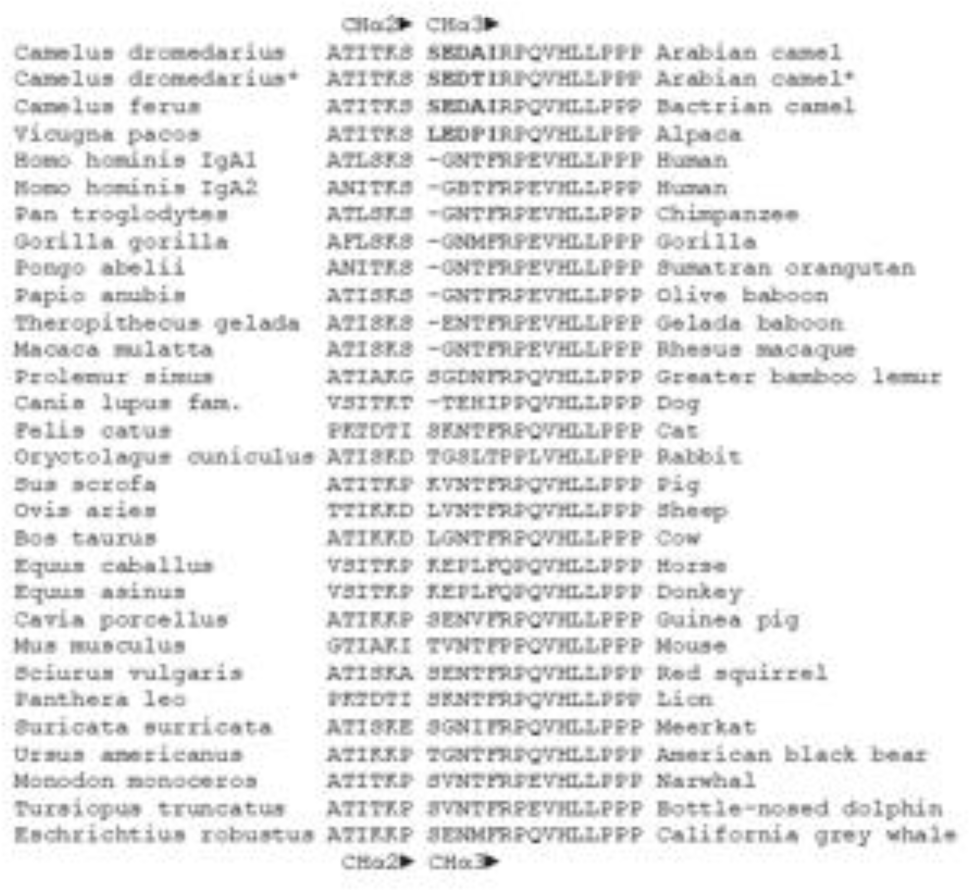
Amino acid sequences of the C_H_α2-C_H_α3 boundary in mammals. Amino acid sequences at the 3’ terminal end of C_H_α2 and the 5’ *terminus* of C_H_α3 domains of the dromedary, Bactrian camel and alpaca were compared with those of other mammals. Names in Latin are to the left. The gap between C_H_α2 and C_H_α3 domains is deliberate. A distinct pentapeptide (bold letters) consisting of serine (leucine in alpaca), glutamic acid, aspartic acid, alanine or threonine (proline in alpaca) and isoleucine was only found in camelids, upstream of a highly conserved arginine followed by an invariant proline at the N-*terminus* of the C_H_α3 domain. Glutamic acid at the 5’ end of C_H_α3 was also present in gelada baboon, horse, donkey, guinea pig, red squirrel, and California grey whale, whereas aspartic acid at this position was identified only in the greater bamboo lemur. In contrast, the most frequently encountered motif in the interdomain region of other mammals at the N-terminal end of the C_H_α3 domain was a tetrapeptide consisting of glycine, asparagine, threonine (or methionine, isoleucine, and valine) and phenylalanine.

### Characterization of IgA heavy chain variants in tissues, milk, and serum by immunoblotting

Proteins of tissue lysates (Fig 5) of the small (lane 1) and large intestine (lane 2), *trachea* (lane 3), third eyelid (lane 4), and also from whey milk (lane 5), dromedary serum (female and male, lane 6 and lane 7, respectively) and human serum from a male donor (lane 8), were separated by SDS-PAGE electrophoresis under reducing conditions, transferred to nitrocellulose membranes and probed with four different polyclonal antibodies raised in rabbits against IgA C_H_ domains: C_H_α1 (panel A), C_H_α2 (panel B), C_H_α3 (panel C) and the IAR (panel D). We predicted that the anti-C_H_α1 antibody would visualize canonical IgA heavy chain, but not the unconventional counterpart, resulting in a single band in immunoblots, whereas anti-C_H_α2, anti-C_H_α3 and anti-IAR antibodies were expected to detect two protein species of different *M*r. As shown in Fig 5A, the anti-C_H_α1 antibody indeed revealed one protein species of estimated *M*r of ∼60 kDa that corresponds to the conventional IgA heavy chain in small (lane 1) and large intestine (lane 2), third eyelid (lane 4), whey milk (lane 5), serum of female (lane 6) and male Arabian camel (lane 7), as well as two bands of ∼60 kDa and ∼65 kDa in human serum, likely cross-reacting with the two known IgA subclasses IgA1 (higher *M*r) and IgA2 (lower *M*r) (lane 8). The signal intensity of canonical IgA heavy chain band was faint in the tissue extract from the *trachea*, reflecting a lower density of plasma cells in the *mucosa* of the upper respiratory tract (lane 3). Surprisingly, the anti-C_H_α1 antibody also detected a polypeptide species with an estimated *M*r of ∼45 kDa in milk and serum, which merits further investigations. In contrast, the expression patterns of IgA heavy chains observed using anti-C_H_α2 (Fig 5B), anti-C_H_α3 (Fig 5C) and anti-IAR (Fig 5D) antibodies, which detect canonical as well as unconventional IgA heavy chain, were similar: the higher *M*r species of ∼60 kDa corresponded to canonical IgA heavy chain, whereas the lower *M*r species of ∼52 kDa represented the unconventional heavy chain variant. Whereas in the tissues examined both IgA heavy chain variants were expressed at comparable levels (lanes 1-4), the ∼60 kDa band was more prominent in milk (lane 5). In circulation, however, only canonical IgA heavy chain was found, favoring the absence of heavy chain-only IgA in the blood (lanes 6 and 7). Weak cross-reactivity was observed with a ∼60 kDa species in human serum (lane 8). Expectedly, the anti-IAR antibody failed to visualize any human IgA heavy chain species as the oligopeptide used for immunization is unique for *Camelidae*. Of note, the band density observed with the anti-IAR antibody appeared identical to that generated by the anti-C_H_α2 antibody, indicating comparable immunogenicity of the 15-mer oligopeptide and the C_H_α2- or the C_H_α3 domains, both comprising ∼100 amino acids.

**Figure 5.**
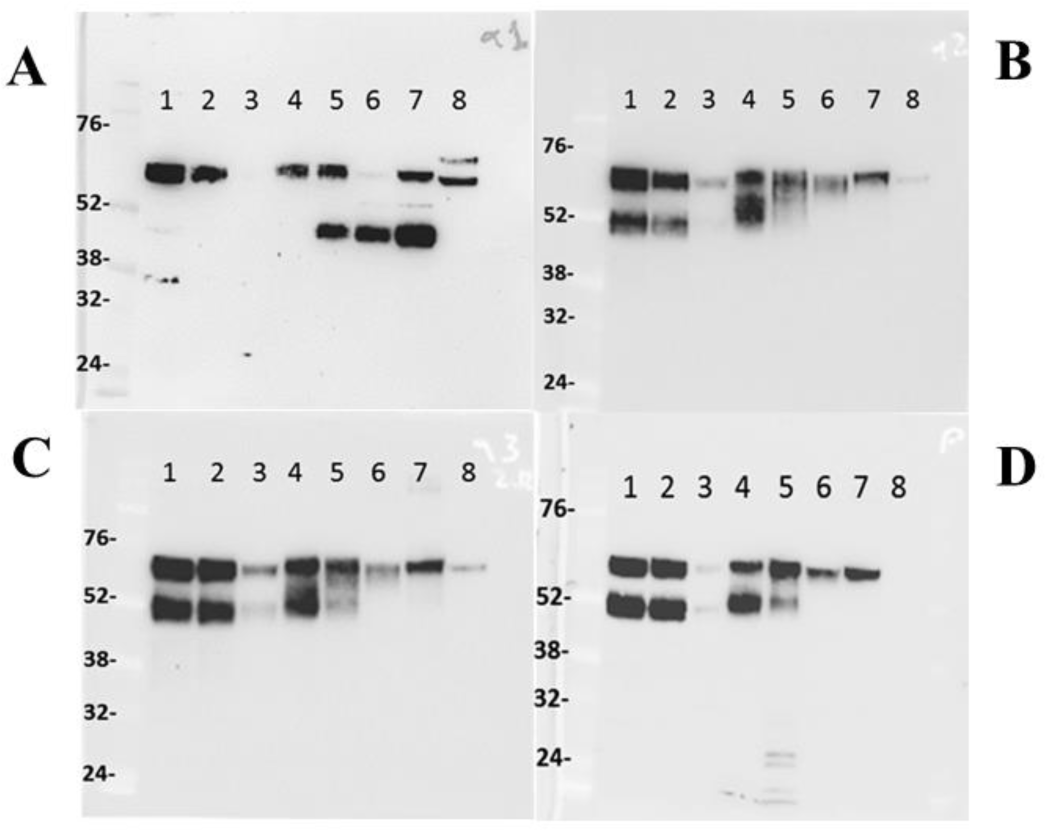
IgA heavy chain variants in the intestine, *trachea*, third eyelid, milk, and serum. One-humped camel proteins (∼20 μg) from small (lane 1) and large intestine (lane 2), *trachea* (lane 3), third eyelid (lane 4), milk (lane 5), female (lane 6) and male (lane 7) serum and human male serum (lane 8) were separated under reducing conditions by 10% (w/v) SDS-PAGE, transferred to nitrocellulose membranes, probed with anti-C_H_α1 (A), anti-C_H_α2 (B), anti-C_H_α3 (C), and anti-IAR (D) antibodies, and visualized with HRP-conjugated anti-rabbit goat antibodies. In (A), the anti-C_H_α1 antibody reacted with a ∼60 kDa species, indicating the expression of canonical IgA heavy chains, with cross-reactivity to two protein species of ∼60 kDa and ∼65 kDa in human serum (lane 8). In camel milk (lane 5) and sera (lanes 6 and 7), an additional prominent protein species of ∼45 kDa was identified by the anti-C_H_α1, but not by other antibodies, most likely representing a proteolytic fragment of canonical IgA heavy chain. In (B), (C) and (D), two protein bands of ∼52 kDa and ∼60 kDa of comparable intensity were identified in tissues (lanes 1, 2, 3 and 4), but not in milk (lane 5) nor serum (lanes 6 and lane 7) where the higher molecular weight species corresponding to canonical IgA heavy chains prevailed. Level of IgA expression in the *trachea* (lane 3) appeared low, likely reflecting lower plasma cell density in the *mucosa* of the upper respiratory tract. Cross-reactivity of circulating human IgA species was observed with all anti-C_H_ domain antibodies, except the camel-specific anti-IAR antibody. Molecular weight markers in kDa are indicated to the left of each panel.

### Expression of IgA heavy chains in MALT by immunohistochemistry

After delineating the histomorphologic features of the third eyelid, we studied by immunohistochemistry the expression of IgA heavy chains not only in the HG but also in other MALT using primary antibodies raised in rabbits against different C_H_α domains (Fig 6-9). The aim was, first, to define the expression patterns of IgA heavy chains in MALT of the dromedary, and second, to determine the spatial distribution of IgA^+^ plasma cells within the tissues examined. We took advantage of purposely generated antibodies specific for C_H_α1, C_H_α2 and IAR, but refrained from using the anti-C_H_α3 antibody on tissue sections because the results obtained by immunoblotting with the anti-C_H_α3 antibody were identical to those observed with the anti-C_H_α2 and anti-IAR antibodies (Fig 5). Based on the results obtained with Western blots we inferred that antibodies binding the C_H_α1 domain will detect classic, yet not unconventional IgA heavy chain, while antibodies binding the C_H_α2 domain and IAR would identify both variants of IgA heavy chain.

**Figure 6.**
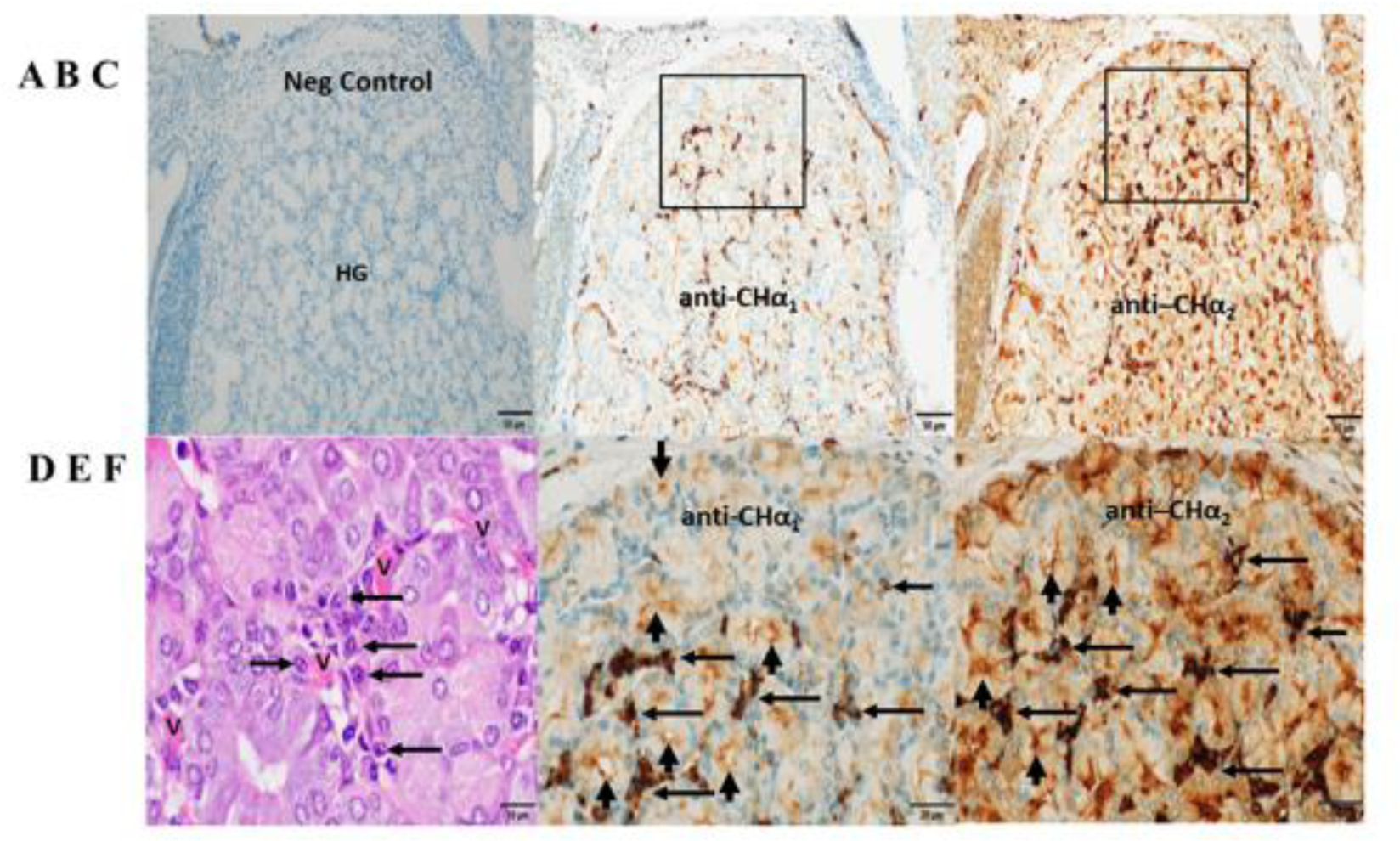
Expression of IgA heavy chain variants in the Harderian gland. Immunohistochemistry was performed on a section through the lacrimal (Harderian) gland (HG) of the third eyelid of the dromedary. Anti-C_H_α1 domain antibodies were used in (B) and (E) and anti-C_H_α2 domain antibodies in (C) and (F) and visualized by goat anti-rabbit HRP conjugated antibodies and hydrogen peroxide/DAB as substrates. The anti-C_H_α1 antibody detected canonical IgA heavy chains, whereas canonical and unconventional IgA heavy chains were both reactive to the anti-C_H_α2 antibody. In (A), no IgA was identified with secondary antibodies only. In (B) and (C), immunoreactivity was observed in the periacinar spaces and within epithelial cells throughout the lobule with anti-C_H_α1 and anti-C_H_α2 antibodies, respectively. In (E) and (F), two rectangular areas selected in (B) and (C) show, at higher power, intense immunostaining primarily in plasma cells in the periacinar spaces (horizontal arrows), and at the luminal apices of epithelial cells (short vertical arrows). In (D), an H&E-stained section demonstrates an agglomerate of typical plasma cells (horizontal arrows) and red blood cells within capillaries (V) adjacent to the basal membranes of *acini*. Magnifications and scale bar lengths were as follows: in (A), (B) and (C), 20x and 50μm, respectively; in (D), 100x and 10 μm, respectively; in (E) and (F), 60x and 20μm, respectively.

**Figure 7.**
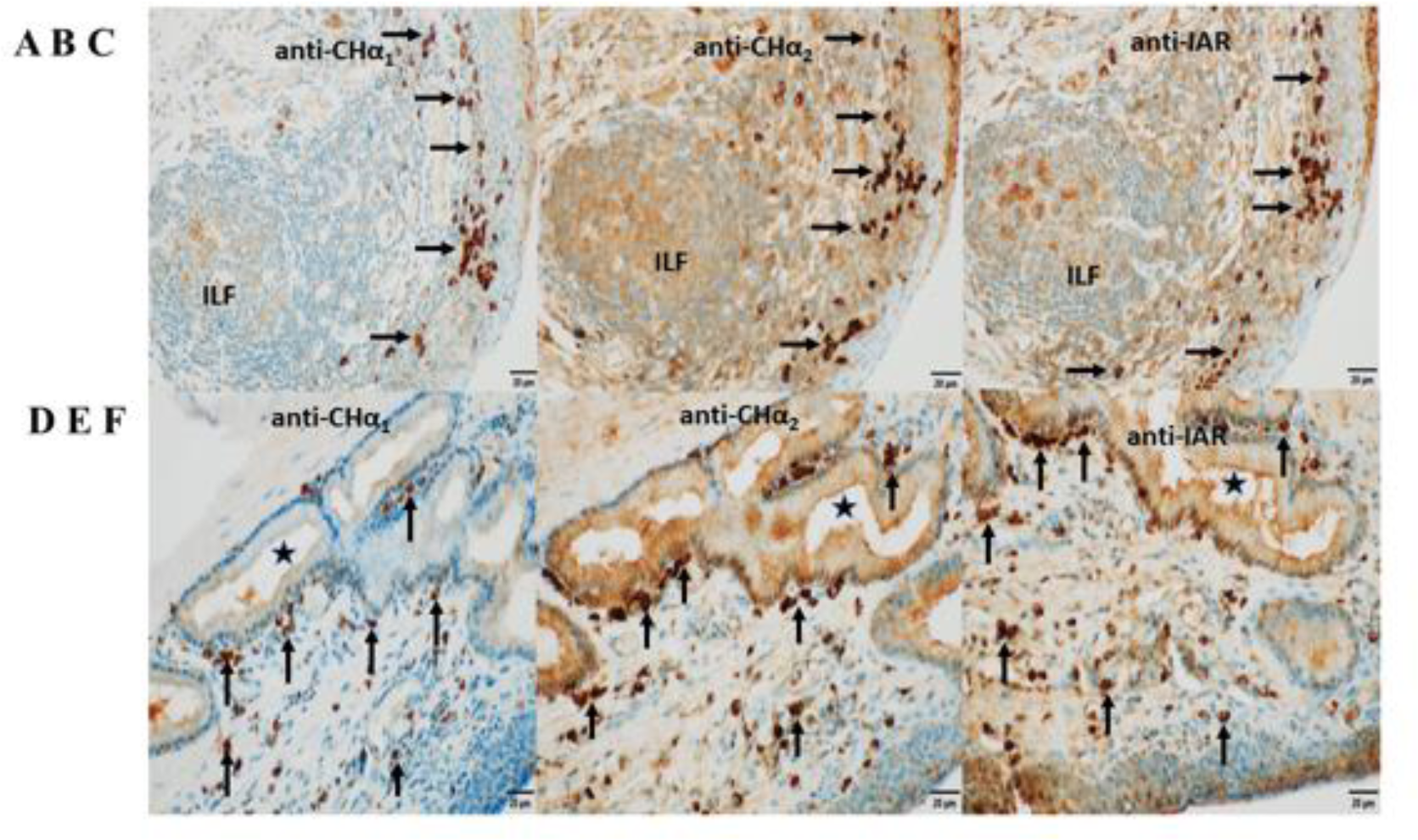
Expression of IgA heavy chain variants in MALT of the third eyelid and *trachea*. Immunostaining was performed on sections through the *conjunctiva* of the third eyelid (top panels) and on sections through the *trachea* (bottom panels) using antibodies raised against C_H_α1 domain in (A) and (D), C_H_α2 domain in (B) and (E), and IAR in (C) and (F), visualized by goat anti-rabbit HRP conjugated antibodies and hydrogen peroxide/DAB as substrates. The anti-C_H_α1 antibody detected canonical IgA heavy chain only. By contrast, both canonical and unconventional IgA heavy chains were identified by the anti-C_H_α2 as well as the anti-IAR antibody. In (A), (B) and (C), IgA heavy chain immunoreactivity to all three antibodies was identified in plasma cells within the stratified squamous epithelial layer of the *conjunctiva* and in the subconjunctival *stroma* (horizontal arrows) as well as within an isolated lymphoid follicle (ILF). IgA heavy chain immunoreactivity was also observed along the surface of the *conjunctiva*. In (D), (E) and (F), IgA^+^ plasma cells (vertical arrows) were noticed alongside the basal membrane of a subepithelial mucous gland lined by columnar *epithelium* and in the subepithelial *stroma* of the tracheal *epithelium*. The *lumen* of the gland is marked by (*). Immunoreactivity was also present within the glandular epithelial layer, reflecting *transcytosis*. Of note, the brown color generated with the anti-C_H_α2 and the anti-IAR antibody was more intense. In this series of microphotographs, magnification and scale bar length was 40x and 20μm, respectively.

**Figure 8.**
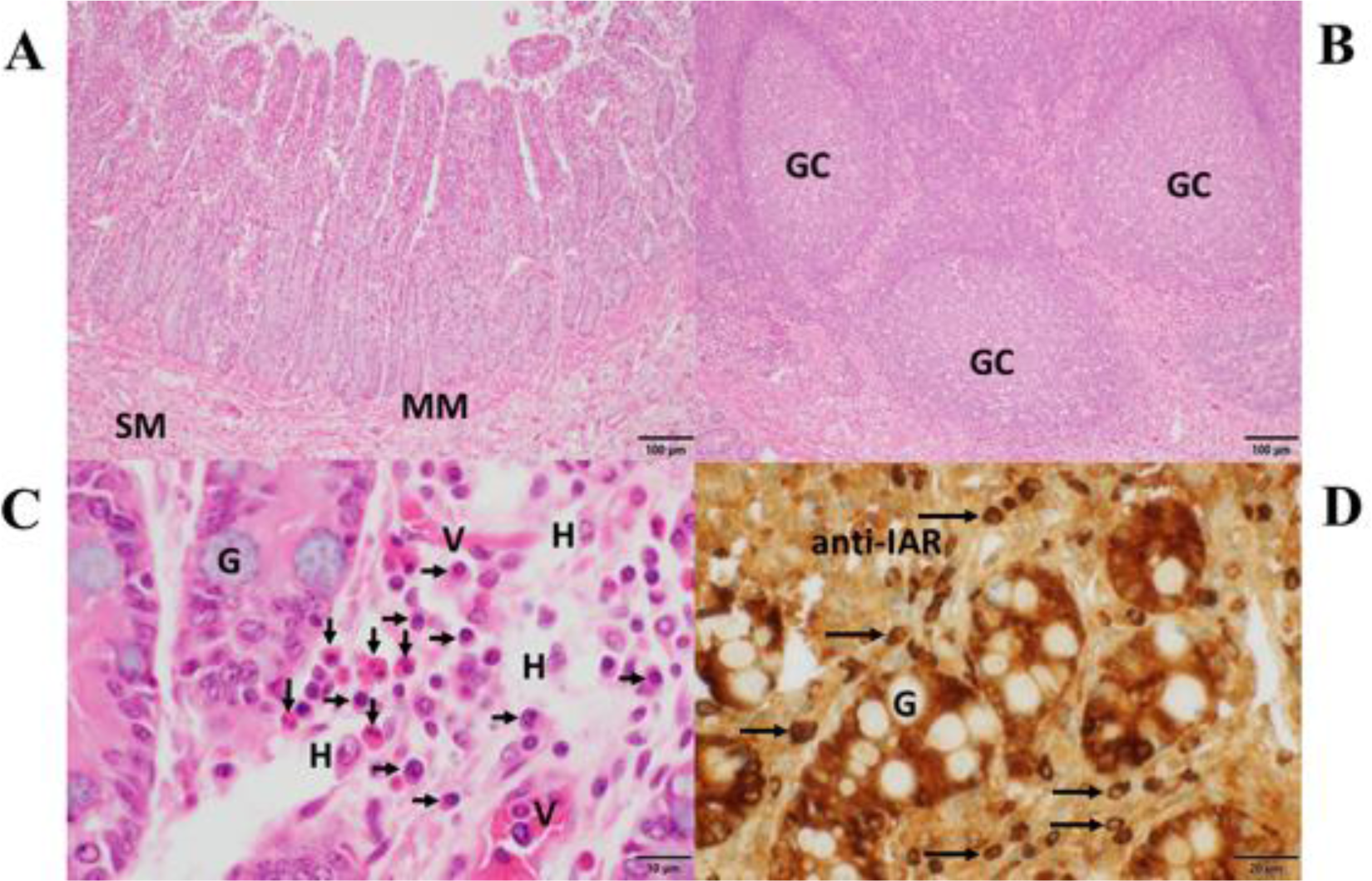
Histomorphology, expression of IgA heavy chains and tissue eosinophilia in the small intestine. In (A), *villi* and crypts characterized the *mucosa* of the small intestine, which was separated from the *submucosa* (SM) by the *muscularis mucosae* (MM). In (B), aggregates of submucosal lymphoid follicles with germinal centers (GC) surrounded by numerous lymphocytes - typical of Peyer’s patches - were present. In (C), details of crypts revealed interspersed goblet cells filled with mucin (G). In the adjacent *lamina propria,* numerous eosinophils (vertical arrows), plasma cells (horizontal arrows), histiocytes (H) and small vessels (V) were observed. Sections in (A), (B), and (C) were stained with H&E. In (D), strong IgA heavy chain immunoreactivity to the anti-IAR antibody was detected in plasma cells (horizontal arrows) and within enterocytes while sparing mucin in goblet cells (G). Magnifications and scale bare lengths were as follows: in (A) and (B), 10x and 100μm, respectively; in (C), 100x and 10μm, respectively; in (D), 60x and 20μm, respectively.

**Figure 9.**
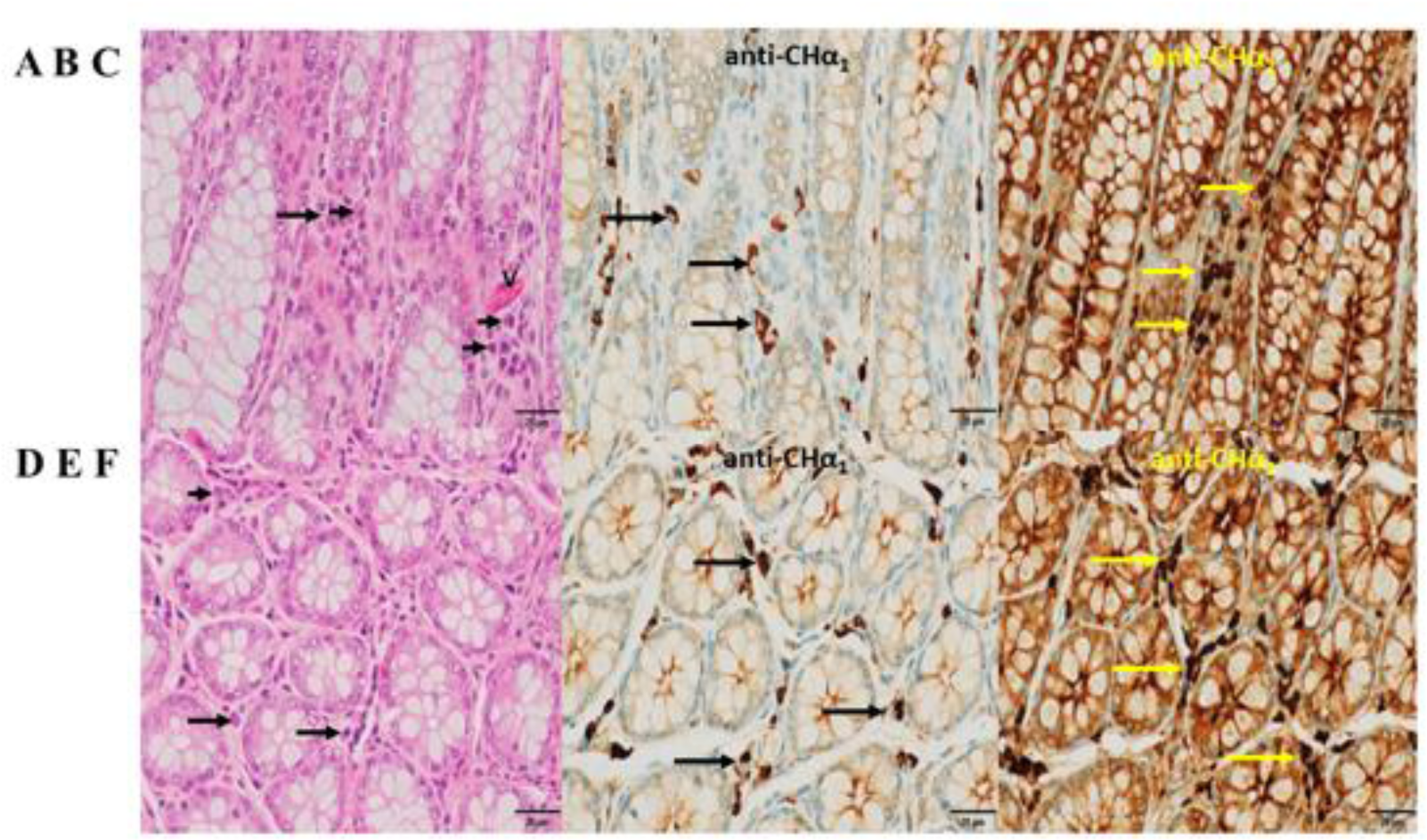
Histomorphology and expression of IgA heavy chain variants in the large intestine. In (A) and (D), the hallmarks of colon *mucosa* are shown in H&E-stained sections. Upper panels depict longitudinal, lower panels transverse sections. Crypts were lined primarily by mucin-enriched epithelial cells (goblet cells) and surrounded by *lamina propria*, where plasma cells (arrows) were found. In (B) and (E), plasma cells showed IgA heavy chain immunoreactivity to the anti-C_H_α1 (arrows), and in (C) and (F) to the anti-C_H_α2 antibody (yellow arrows). Immunoreactivity was also observed in the epithelial cells within crypts, indicative of *transcytosis*. In (A), a capillary (V) was identified by the presence of red blood cells. In contrast to the *lamina propria* of the small intestine (Fig 8C) eosinophils were absent in this area of the colon. In all microphotographs magnification was 60x, and scale bar length was 20μm.

In sections through a lobule of the HG (Fig 6), we noticed that classic IgA heavy chain was expressed in plasma cells throughout the HG using the anti-C_H_α1 antibody, as shown in Fig 6B, and at higher magnification in Fig 6E. We surmise that heavy chains of both IgA subclasses were co-expressed in plasma cells using the anti-C_H_α2 antibody, as shown in Fig 6C and at higher magnification in Fig 6F. IgA^+^ plasma cells are indicated by horizontal arrows (Fig 6E, F). A negative control section is shown (Fig 6A), and, in an H&E-stained section (Fig 6D), a nest of plasma cells displayed the characteristic morphology, with eccentric nucleus, condensed chromatin resulting in “clock face” morphology and cytoplasmic “Hof” (horizontal arrows). These plasma cells resided in the periacinar *stroma* adjacent to the basal lamina of the columnar epithelial cells and next to capillaries (V) identified by intravascular red blood cells. The more intense brown color observed with both, the anti-C_H_α2 antibody in Fig 6, 7 and 9, as well as the anti-IAR antibody in Fig 7 and 8, was possibly related to non-specific background staining seen throughout the section. However, we hypothesize that an additive effect of immunoreactivity resulted in the apparent darker brown color as anti-C_H_α2 and anti-IAR antibodies react to both types of IgA heavy chains, whereas the anti-C_H_α1 antibody was selective for canonical IgA heavy chains. Immunostaining was also observed at the apical pole of acinar epithelial cells towards the luminal surface (Fig 6E, F; short vertical arrows), where IgA is transported *via* polymeric Ig receptor (pIgR)/secretory component (SC)-mediated *transcytosis* (49).

In CALT (Fig 7, top row), plasma cells (horizontal arrows) expressing IgA heavy chains detected by anti-C_H_α1 (Fig 7A), anti-C_H_α2 (Fig 7B) and anti-IAR (Fig 7C) antibodies are shown. They resided along the basal lamina of the stratified *conjunctiva* on the bulbar side of the third eyelid overlying an ILF, while some of them were found within the *epithelium*. The observation of intraepithelial plasma cells was also reported in other *Artiodactyla*, horse as well as the duck (22, 50). ILFs showed weak central immunoreactivity compatible with intrafollicular IgA class switching. Bottom panels in Fig 7 demonstrate IgA^+^ plasma cells (vertical arrows) reactive to anti-C_H_α1 (Fig 7D), anti-C_H_α2 (Fig 7E) and anti-IAR (Fig 7F) antibodies in the subepithelial *stroma* of the pseudostratified columnar *epithelium* of the *trachea* and in the vicinity of a subepithelial mucous gland lined by columnar *epithelium* (*). We attribute the immunostaining of epithelial cells, which was more intense with the anti-C_H_α2 and the anti-IAR antibody, to the *transcytosis* of IgA. Of note, the intensity of the color reaction with the anti-IAR domain antibody appeared identical to that generated by the anti-C_H_α2 antibody suggesting comparable immunogenicity or avidity.

The histomorphology of the *mucosa* of the small intestine is shown in Fig 8. V*illi* and crypts are separated by the *muscularis mucosae* (MM) from the *submucosa* (SM) (Fig 8A). Aggregations of lymphoid follicles with germinal centers (GC), typical of Peyer’s patches are shown in Fig 8B. Numerous eosinophilic neutrophils (vertical arrows), abundant plasma cells (horizontal arrows) along with scattered histiocytes (H) were the principal cellular components in the highly vascular (V) *lamina propria* (Fig 8C). A few goblet cells (G) were found between the enterocytes. In Fig 8D, IgA heavy chains expressed in plasma cells (horizontal arrows) and in enterocytes exhibited strong reactivity to the anti-IAR antibody, whereas the mucin in goblet cells (G) remained unstained.

In the *mucosa* of the large intestine (Fig 9), with crypts lined by numerous goblet cells, plasma cells (horizontal arrows) were found in the *lamina propria* between crypts as shown in longitudinal (upper panels) and transverse sections (lower panels). In Fig 9B and 9E, representative plasma cells (horizontal arrows) showed IgA immunoreactivity to the anti-C_H_α1 antibody, and in Fig 9C and 9F, to the anti-C_H_α2 antibody. Immunostaining was also associated with colonic epithelial cells, indicative of IgA *transcytosis*, but not with mucin in goblet cells. In contrast to the small intestine, eosinophils were absent in this section of the large intestine.

## Discussion

We demonstrate that dromedary IgA, the protagonist immunoglobulin class at mucosal sites in mammals, maintains the distinctive dualistic structure of canonical and unconventional heavy chains, which was discovered in the IgG class of the one-humped camel and later in the nurse shark (1, 4). We also show that classic and unconventional IgA heavy chains are co-expressed in MALT of the Arabian camel.

### The third eyelid harbors key components of conjunctival immune defense

The nictitating membrane was chosen as a relatively easily accessible source of IgA because it integrates inductive and effector components of mucosal immune defense against airborne antigens. The histomorphology of the third eyelid was extensively explored and compared in many quadrupeds, including dromedaries and alpacas (23, 51), and the sites of synthesis of IgA and SC determined (22). On the immune-inductive site, CALT consists of characteristic mucosal ILF lacking germinal centers located underneath the innermost (bulbar) aspect of the nictitating membrane, whereas at immune-effector sites, plasma cells sparse throughout the *subconjunctiva* or clustered in the periacinar *interstitium* of the HG express IgA heavy chains that engage *via* the J-chain the epithelial pIgR located at the basolateral surface of the glandular epithelial cell for *transcytosis*, eventually resulting in the release of sIgA, a multimeric protein complex that includes the SC, which is proteolytically generated from the pIgR (52, 53). Based on data obtained from analyses of *nasopharynx*-associated lymphoid tissue situated in the tonsils and gut-associated lymphoid tissue, both representing inductive sites for B cell differentiation, it can be inferred that in the ILF of CALT, similar immunoregulatory events of B cell maturation, including IgA class-switching, take place (54–56). ILF located in the *lamina propria* below the *epithelium* of the third eyelid’s bulbar *conjunctiva*, constitute an immune-inductive site, whereas IgA^+^ plasma cells residing in the *lamina propria* of the *conjunctiva* or grouped in the periacinar *stroma* of the HG constitute the effector arm of conjunctival humoral immunity. From inductive sites, IgA^+^ B cells are thought to migrate *via* the lymphatic system to reach regional lymph nodes, where they acquire redirecting homing molecules for regionalized tissue-specific localization in the *lamina propria* of the *conjunctiva* and in the periacinar *stroma* of the lacrimal gland. However, given the proximity of inductive and effector sites in the third eyelid, it is tempting to speculate on an alternative, T-cell independent pathway in which IgA^+^ B cells bypass further maturation in lymph nodes and directly access their terminal effector niches (57).

### Unconventional and classic IgA heavy chains are expressed side-by-side in MALT

The RACE results showed heterogeneity in the length of the 5’ PCR products in contrast to a single 3’ PCR species. Sequencing of two 5’ PCR species running at ∼1100 bp confirmed that they were conventional IgA heavy chains encompassing the ∼300 bp C_H_α1 domain, whereas most clones with the ∼800 bp 5’ PCR products were void of the C_H_α1 domain, demonstrating that plasma cells at the effector site of the third eyelid were capable of co-expressing both types of IgA heavy chains. 119 deduced amino acid sequences of V_H_H and constant domains from our IgA cDNA library are listed in the Supplementary file#1. The 19-mer leader oligopeptide, MELGLSWVVLAALLQGVQA, was earlier found associated with variable domains of classic (V_H_) as well as unconventional heavy chains (V_H_H) in a genomic library generated from dromedary liver DNA (58). The V domains following this particular leader sequence derive from a subgroup of *IGHV3* genes (equivalent to human *IGHV3* family clan III) (59). We found that the cardinal amino acid substitutions in the V_H_H domain of unconventional IgA heavy chain were identical to those reported for the V_H_H domain of IgG (2). These substitutions were as follows: in FR1, serine for leucine; in FR2, phenylalanine for valine, glutamic acid for glycine, arginine for leucine, and glycine for tryptophan. It was reported that these amino acids reshape the protein surface and increase the solubility of isolated V_H_H domains (60, 61). Regarding the length of CDR in the V_H_H domain, the rearranged CDR3 is more extended than in other species with a maximum of 26 amino acids in our IgA heavy chain cDNA library. However, there was considerable length variation among the CDR3, less so in CDR2, as shown in Supplementary file#1. Of interest was the presence of a short hinge region in all clones, which consisted of six consecutive prolines, preceded by a valine, almost identical to the proline pentamer of human IgA2 and chimpanzee IgA2 hinges, known to confer remarkable resistance to highly specific IgA1 bacterial proteases (62).

Co-expression of both IgA heavy chain variants in tissue extracts was not only observed in the third eyelid, but also in the *trachea*, as well as in the upper and lower intestinal tract, and inferred by immunohistochemistry in plasma cells of the HG, the *conjunctiva*, *trachea*, and upper as well as lower intestine. The findings were different in milk, where classic IgA heavy chain prevailed, and in serum, where unconventional heavy chain-only IgA was undetectable. The latter observation contrasted the original discovery of circulating heavy chain-only IgG in the serum of dromedaries which amount to ∼3/4 of total IgG (1). However, the apparent absence of unconventional IgA heavy chain in serum could be accounted for by very low IgA levels, analogous to the situation in human blood where IgA levels reach ∼1/5 of the concentration of IgG, and therefore very low levels of the heavy chain-only variant could escape detection. Of interest was the demonstration of the presence of a ∼45 kDa protein species in circulation, which was observed with the anti-C_H_α1 antibody only. It may represent a cleaved fragment of classic IgA since serum and milk contain a variety of proteolytic systems that generate Fab fragments of IgA (63) and IgG class as well (64, 65). Bacterial IgA1 proteases exist that cleave proline-serine bonds specifically in the IgA1 hinge region (66), and two proline-serine bonds are found upstream and central in the C_H_α2 domain of the dromedary. If enzymatic cleavage at these sites with subsequent degradation of the C-terminal fragment took place, the non-reactivity of the anti-C_H_α2, anti-C_H_α3 and anti-IAR antibody would find an explanation. Cross-reactivity of the anti-C_H_α1 antibody with another immunoglobulin class circulating in serum or secreted in the milk offers an alternative, yet less likely, explanation. Further characterization of the observed protein species is needed.

### A cryptic acidic oligopeptide at the 5’ *terminus* of the C_H_α3 domain distinguishes camelids from other mammals

At the N-*terminus* of the C_H_α3 domain, we noticed a novel pentapeptide, consisting of serine, glutamic acid, aspartic acid, alanine or sporadically threonine, and isoleucine. This sequence was identical in the Bactrian camel, whereas we found threonine for alanine also in the dromedary IgA* classic heavy chain. At the N-*terminus* of the C_H_α3 domain in alpaca, leucine was observed instead of serine, and proline for alanine. A comparison with other mammals revealed that only camelids have a pair of acidic amino acids and isoleucine at this site of transition between the C_H_α2 and C_H_α3 domains, where immediately 3’ to the pentapeptide a highly conserved arginine followed by an invariant proline marks the N-*terminus* of the C_H_α3 domain (47, 48). However, two amino acid sequences (No 109 and No 111 in the Supplementary file) were an exception to this rule as the pentapeptide was found to contain not acidic but basic residues (arginine and histidine). By contrast, in other mammals a tetrapeptide consisting of glycine, asparagine, threonine, and phenylalanine represents the most common motif in the interdomain region, with the notable exception of the dog (threonine-glutamic acid-histidine-isoleucine), rabbit (glycine-serine-leucine-threonine), horse and donkey (glutamic acid-proline-leucine-phenylalanine), and greater bamboo lemur (glycine-aspartic acid-asparagine-phenylalanine). We surmise that in camelids, and possibly a few other species, negatively (or rarely positively) charged amino acids centered between basic residues, *i.e.* lysine upstream, arginine and histidine downstream, may affect structural and functional properties of the interdomain region likely accounting for novel protein-protein interactions with a putative receptor or a soluble protein.

### Previous studies on MALT in camelids failed to investigate the co-expression of heavy chain-only and conventional IgA or IgG

Reports on the mucosal immune system in camelids in general and on the IgA system in particular are scarce. Recently, the topography of the pIgR was described in the lung of the Bactrian camel (67). Earlier, the small intestine was thoroughly examined in Bactrian camels where the concurrent expression of IgA and IgG was found and interpreted as a functional dual barrier of mucosal immune defense (68, 69). Predominant IgA over IgG expression was also described in a specific area of the three-chambered stomach of the dromedary (70). Yet, the probing antibodies used to locate these isotypes were not designed to differentiate between canonical and heavy chain-only antibodies and molecular data were not provided. Therefore, the question whether heavy chain-only antibodies existed in the gut of the Bactrian camel was not answered. Based on our observations an additional defense line of mucosal adaptive immunity in the dromedary must now be considered since both variants of IgA heavy chains were jointly and comparably expressed as judged by the density of the protein bands obtained by immunoblotting. In addition, IgA^+^ plasma cells were detected at effector sites in the *lamina propria* of the *mucosa* of the small as well as the large intestine by anti-C_H_α1, anti-C_H_α2 and anti-IAR antibodies, with the first antibody selective for conventional IgA heavy chains, while the other two antibodies were reactive to both IgA heavy chain variants. Although the darker brown color observed with the anti-C_H_α2 and anti-IAR antibodies compared to the staining intensity of the anti-C_H_α1 antibody suggested an additive effect, we cannot exclude that the differences in color intensities were caused by non-specific background staining. We were therefore unable to determine the ratio of IgA^+^ plasma cells producing classic heavy chains versus those expressing unconventional heavy chains. Yet, the results obtained from Western blots suggest that they were in *equilibrium* at the time the tissue samples were obtained. Moreover, we assume that in the dromedary plasma cells have the potential to switch transcription between either IgA heavy chain variant, an unlikely scenario in the gut of the Bactrian camel or the alpaca which possess only one *Cα* gene (71, 72).

### Potential interactions of plasma cells and eosinophils in the intestinal mucosa

Another observation deserves attention, namely the proximity of eosinophils and plasma cells in the *lamina propria* of the small intestine, but not in a sample of the lower bowel, suggesting a functional relationship between terminally differentiated cells of innate and adaptive immunity. We do not know whether in the small intestine of the Bactrian camel similar conspicuous eosinophilic infiltrates exist in proximity of plasma cells. The role of eosinophils in providing key survival factors, such as APRIL (a proliferation-inducing ligand), IL-6, IL-4, IL-10 and TNF-α, to plasma cells has been described in the bone marrow, whereas in the digestive tract of mice, eosinophils maintain the function of IgA^+^ plasma cells and participate in T cell-independent class-switching *via* a TGF-β-dependent mechanism (73–76). Therefore, it is conceivable that eosinophils, beside their pivotal effector function in innate and FcεR-mediated immune responses against intestinal parasites, *e.g. trichuris* and *nematodirus* species, modulate the synthesis of IgA heavy chains and promote survival of IgA^+^ plasma cells in niches of the intestinal *lamina propria* in the dromedary (77–79). Furthermore, interactions between eosinophils and IgA^+^ plasma cells may be mutual as IgA binds to and activates eosinophils via a highly glycosylated isoform of FcαRI (80).

### Nucleotide and protein sequence databases of IgA in camelids were not in support of side-by-side expression of IgA heavy chain variants in the dromedary

The existence of heavy chain-only IgA in the dromedary is at odds with available data in other members of the *Camelidae* family, the closest species being the two-humped Bactrian camel and the more distantly related South American family member alpaca, which both miss a second *Cα* gene, a *sine qua non* for the transcription of unconventional Ig heavy chains. In the *IgH* locus of the alpaca, only one gene for *Cα, Cδ, Cε* and *Cμ* was identified in contrast to four *Cγ* genes, whereas Bactrian camels possess six *Cγ* genes and one gene for *Cα, Cδ, Cε* and *Cμ* (71, 72, 81). Moreover, partial protein sequences of IgA deduced from a genomic library in Bactrian camel (GenBank EPY85672.1) (71) and alpaca (GenBank CAO795581.1) (72) as well as from a spleen cDNA library of the Bactrian camel (GenBank KP999944.1) also contain the C_H_α1 domain, indicating that IgA in these *Camelidae* is composed of a *bona fide* IgA heavy chain. While five *Cγ* genes were found in the dromedary, conclusive numbers on *Cα* genes were not revealed (10). Whether our findings of the co-expression of canonical and unconventional IgA heavy chains can be extrapolated to all dromedaries demands confirmation as phenotypically different breeds of dromedaries exist, 14 alone in Saudi Arabia (6, 82, 83). Therefore, we cannot at present generalize these findings because the genetic makeup could differ among various breeds (84). The dromedaries examined here belonged to the Hadhana breed, as judged by the light brown color of their coat. Given the numerical variation of *Cγ* genes in *Camelidae* (*i.e.* five in dromedary, six in Bactrian camel and llama, four in alpaca), the presence of a second *Cα* gene in one-humped camels coding for an alternative IgA heavy chain is plausible. Two genes code for IgA subclasses, IgA1 and IgA2, in humans, chimpanzees and gorillas, and gibbons express two IgA subclasses as well, whereas most other species examined have just one *Cα* gene (85–87). Concerning the numbers of *Cα* genes, however, the absolute champion is the rabbit, with a remarkable number of 13 *Cα* genes (88).

Hypothesis: were duplication and intronic mutation of the *Cα* gene confirmed by genomic analyses in other breeds of dromedaries we speculate that this event occurred late in evolution when one-humped camels emerged and migrated to the deserts of the Indian subcontinent, Southwest Asia, Arabian Peninsula, North and East Africa

Considering the advances in characterizing the mucosal immune system in general and IgA in particular (89–91) with its eminent role in natural and induced mucosal immune defense against newly emerged viral pathogens such as SARS-CoV-2 in humans (92–95) and MERS-CoV in the dromedary (96, 97), it was our aim to determine whether in the dromedary both subclasses of IgA antibodies co-exist at mucosal surfaces. Our study demonstrates that the IgA class maintains the typical, camelid-specific dualism of antibody structure which was acquired over millions of years throughout transcontinental migration granting the control of infections under the harshest climatic and extreme physiologic conditions (98). There is evidence that the affinity of heavy chain-only antibodies of the IgG class for their epitopes is superior to that of canonical tetrameric antibodies, providing a clear advantage in antigen recognition and neutralization, and the same could be translated to the IgA class (11). A conclusive explanation, however, why functional antibodies with a simpler structure emerged in two evolutionary remote, terrestrial (*Camelidae*) and aquatic creatures (nurse shark), remains so far elusive. Furthermore, the detection of canonical and unconventional IgA heavy chain in dromedaries adds another firewall of immune defense at mucosal surfaces as other species of the *Camelidae* family do not possess a second *Cα* gene at the heavy chain locus with the specific G → A transition mutation in the canonical splicing site, *i.e.* GT, flanking the first exon, which by analogy with *Cγ* genes, is the *sine qua non* for transcribing unconventional heavy chain genes (3, 99, 100). Our findings expand the antimicrobial spectrum of single-domain heavy chain-only antibodies to the mucosal membranes of the eye, the upper respiratory tract, and the intestine. Genomic data will determine whether in the dromedary the expression of the unconventional heavy chain-only IgA variant is indeed the result of a gene duplication event combined with a splicing defect that occurred when one-humped camels emerged, and whether there are further IgA subclasses or allotypes.

## Acknowledgements

We thank Dr A. Alaiya, Dr R. Al Hijailan, Mrs A. Inglis, Mrs A. Alanazi, Mrs M. Kafaar and Mr I. Keragala for technical assistance and Mrs C. Touzinte and Mr M. Noreldin for secretarial work.

## Funding Statement

This work was funded by grant Nr. 11-BIO2073-20 (The Long-Term Comprehensive National Science, Technology and Innovation Plan) and grant Nr. 42-1 (MERS Research Grant Program) from King Abdulaziz City for Science and Technology (KACST), Riyadh, Saudi Arabia. UK is supported by an UAEU grant.

## Abbreviations/Acronyms

Amino acids: one-letter symbols
bp: base pair
CALT: *conjunctiva*-associated lymphoid tissue
CDR: complementarity determining region
C: constant
C_H_: constant heavy
DAB: 3,3’-diaminobenzidine tetrahydrochloride
FR: framework region
GSP: gene specific primer
HC: heavy chain
H&E: hematoxylin and eosin
HG: Harderian gland
HRP: horseradish peroxidase
IAR: inter-α region
Ig: immunoglobulin
ILF: isolated lymphoid follicle
MALT: *mucosa*-associated lymphoid tissue
PCR: polymerase chain reaction
pIgR: polymeric Ig receptor
RACE: rapid amplification of cDNA ends
SC: secretory component
SG: sebaceous gland
sIgA: secretory IgA
Tp: tailpiece
V: variable
V_H_: variable heavy
V_H_H: variable heavy heavy

## Legend of Supplementary file#1

### Primary structure of unconventional IgA heavy chains in the dromedary

119 amino acid sequences of dromedary unconventional IgA heavy chain, deduced from their corresponding cDNA templates originating from the nictitating membrane, were aligned according to IMGT^®^ delimitations. Amino acid residues are marked with different colors. The *bona fide* immunoglobulin fold of the variable domain consists of 9 antiparallel β strands (A, B, C, C’, C’’, D, E, F, G), which include four framework regions (FR1, FR2, FR3 and FR4), and three adjacent loops, *i.e.* BC-loop, C’C’’-loop, and FG-loop, that correspond to complementarity determining regions CDR1, CDR2 and CDR3, respectively (46). This distinctive structure is preserved in the variable domain of dromedary heavy chain-only IgA, termed the variable heavy heavy (V_H_H) domain. A conserved 19-mer leader peptide, MELGMSWVVLAALLQGVQA (MELG leader peptide), precedes the V_H_H domain, consisting of FR1, followed by CDR1, FR2, CDR2, FR3, CDR3 and FR4, which was separated from the C_H_α2 domain by a short, proline-rich hinge region (VPPPPPP). The V_H_H domains derive from a set of dromedary *IGHV3* genes (equivalent to human *IGHV3* family clan III). In our library, the highest frequency of *IGHV3* genes in the V-D-J rearranged CDR3 was found for *IGHV3-23*04* and *IGHV3-66*02*, followed by *IGHV3-74*02*. The multi-colored, mosaic-like appearance of CDR3, combined with length differences ranging from 14 to 26 amino acids (each dash indicates the absence of a residue), mirrors the typical hypervariability of antigen contact sites. C-terminal to CDR3, starting with FR4, followed by the proline-rich hinge region, constant domains C_H_α2 and C_H_α3, and ending with the invariant 18-amino-acid tailpiece, the variability of amino acid sequences abruptly ceases, marking the boundary between variable and constant domains. FR4 encompasses the C-terminal part of the *J* gene segment, with the canonical J motif W-G-X-G. In the V_H_H domain, unique, camelid-specific amino acid substitutions (phenylalanine for valine, glutamic acid for glycine, arginine for leucine, glycine for tryptophan) are found in FR2 (C strand and C’ strand), and serine replaced leucine in FR1 (A strand). Beside the exceptional length variability of CDR3 and the absence of C_H_α1, these substitutions represent the fingerprint of the V_H_H domain in heavy chain-only antibodies. The camelid-specific amino acid changes from hydrophobic to hydrophilic residues are positioned in the GFCC’C’’ β sheet of the immunoglobulin fold, which in the classic tetrameric configuration forms the V_H_-V_L_ interface, superfluous in the heavy chain-only variant (2). Apart from two invariant cysteines in FR1 (B strand) and FR3 (F strand), additional cysteines in CDR1 and CDR3 stabilize the distinct conformation of the paratope in heavy chain-only antibodies. Of note is the distinct amino acid composition of the 5’ end of the C_H_α3 domain, consisting of two anionic residues (E and D) in most of the IgA heavy chain-only sequences, except for sequence No 109 and No 111, where basic residues (R and H) were observed instead. The polar character of the N-terminal region of the C_H_α3 domain may account for a hitherto unexplored protein-protein interaction.

### Contribution to the field statement

Camelids share a unique feature of antibody structure, namely the existence side-by-side of homodimeric antibodies consisting of a pair of two heavy chains void of light chains (heavy chain-only antibodies) and classic tetrameric antibodies composed of two heavy and two light chains. The dichotomy was first discovered in the IgG class of the dromedary, and later confirmed in two-humped Bactrian camels, as well as in the New World camelids lama and alpaca. The dualism of antibody structure in camelids resulted from gene duplication events combined with a mutation at the canonical splicing site located at the 3’ C_H_1/intron border. Although this phenomenon was not reported in other Ig classes in camelids, definitive data on IgA in the dromedary were lacking. In this study, which was carried out on the third eyelid, as well as the upper respiratory and intestinal *mucosae* of the one-humped Arabian camel, we show that, in contrast to other species of the *Tylopoda* suborder, IgA is indeed composed of heavy chain-only and conventional antibodies, thereby conferring the apparently superior antimicrobial potency of unconventional, single-domain, heavy chain-only antibodies on the mucosal surfaces of the *conjunctiva*, the upper respiratory tract and the gut.

---

**Supplementary Figure 1.**
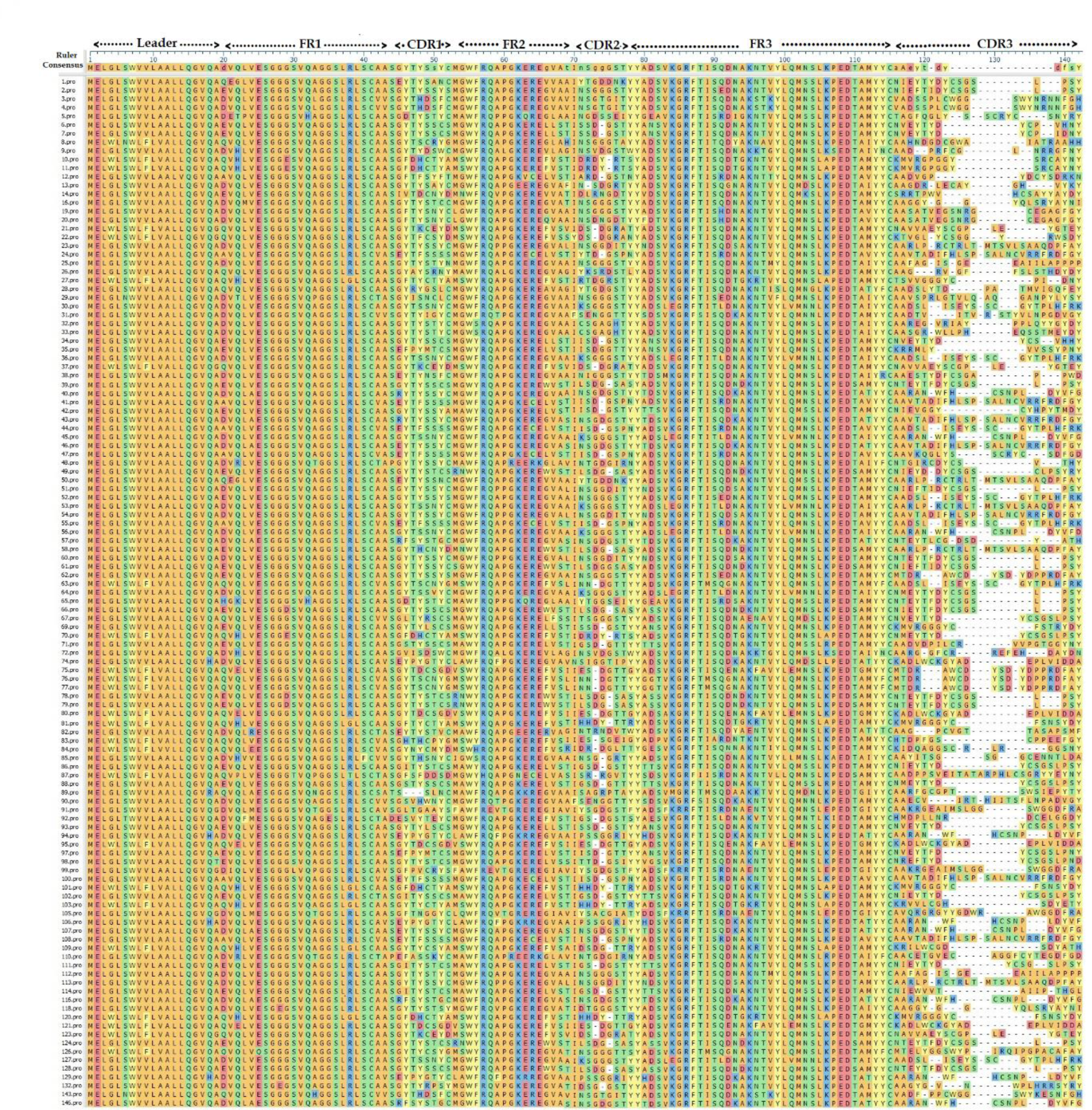

**Supplementary Figure 2.**
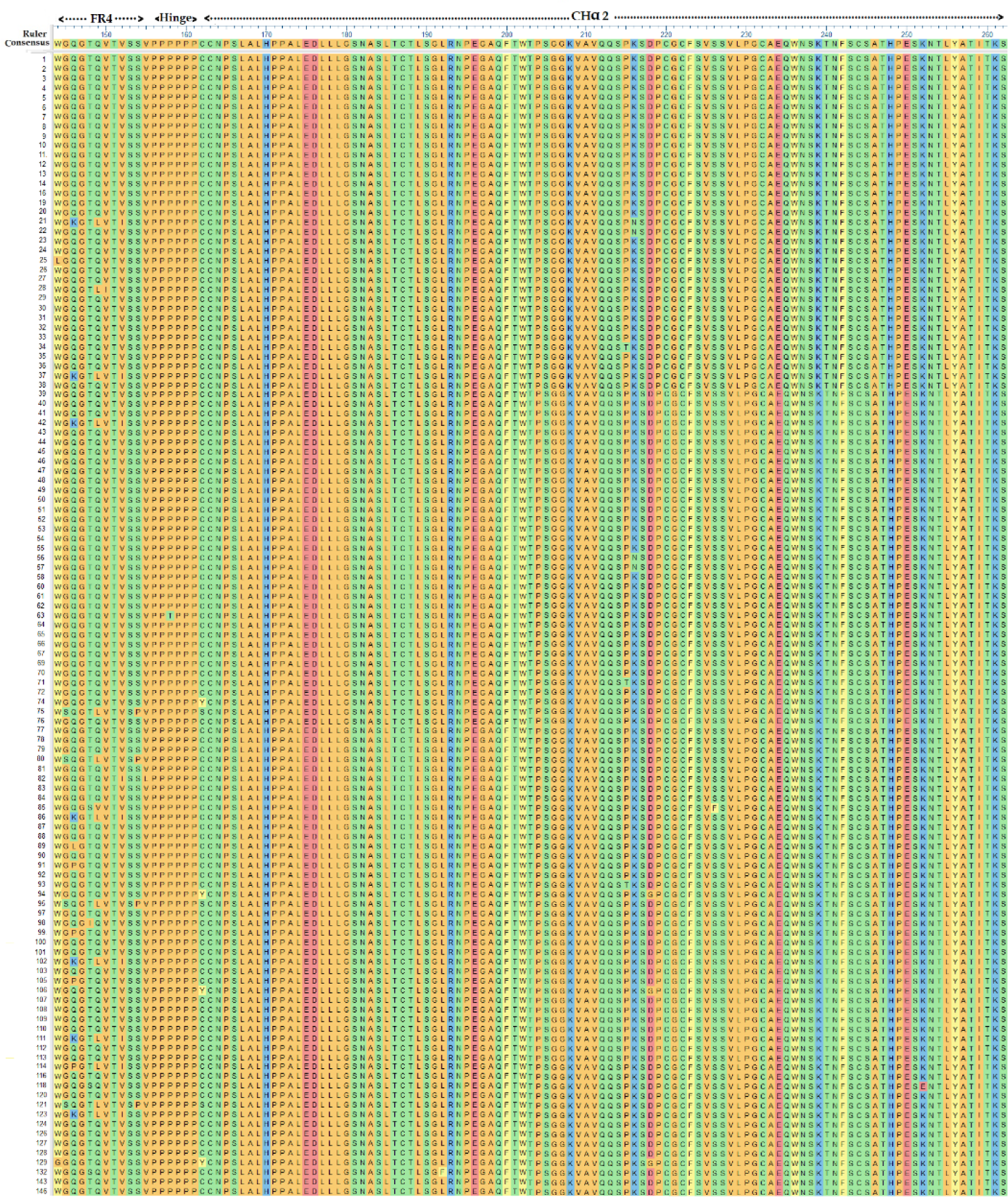

**Supplementary Figure 3.**
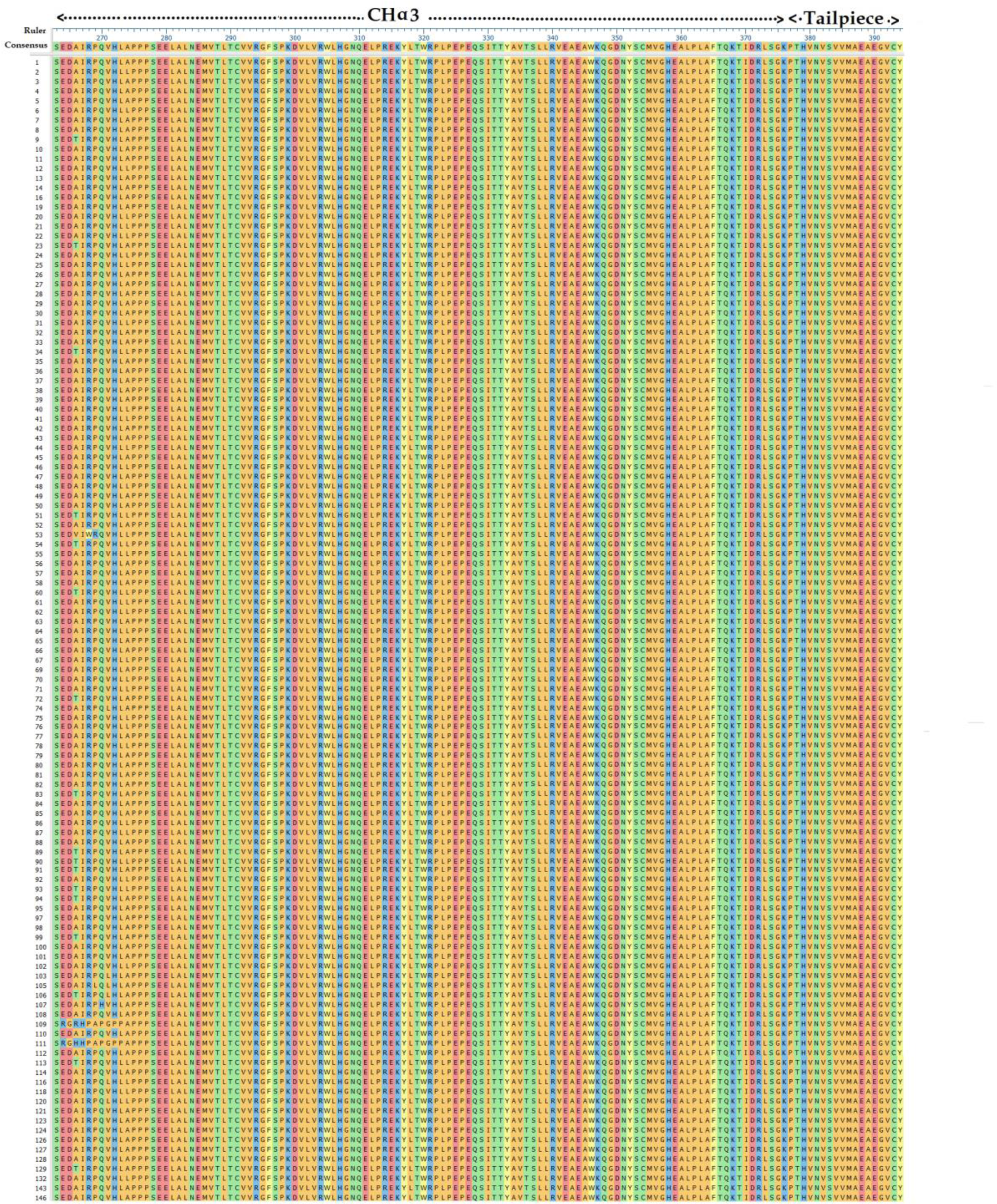

## Notes

### Competing Interest Statement

The authors have declared no competing interest.

